# OmniGene-4: A Unified Bio-Language MoE Model with Router-Level Interpretability and Modality-Invariant Transfer

**DOI:** 10.64898/2026.05.12.724542

**Authors:** Liang Wang

## Abstract

How do multi-modal large language models that jointly process natural language and biological sequences (DNA, protein, structural alphabets) actually answer biological questions, especially sequence-grounded questions whose answer depends on residue-level patterns rather than literature recall? We introduce **OmniGene-4**, a unified bio-language Mixture-of-Experts foundation model on Gemma-4-26B-A4B (128 experts/layer, top-8 routing), and use its discrete router state to dissect this question. By hooking every router across eight task families, we provide the first router-level decomposition for a biological MoE: continued pretraining (CPT) accounts for **96%** of cross-task expert differentiation and supervised fine-tuning (SFT) for **4%**, reshaping middle and output layers respectively. Within the protein-homology task family, per-pair routing divergence stays below 0.04 (vs 0.23 cross-task), implying that sequence-grounded decisions occur *inside* expert computation rather than at the gate — the gate selects the modality, the experts compute the answer. The pipeline yields strong benchmarks: remote-homology **82.60%** (vs ESM-2 3B, MMseqs2, DIAMOND by 28–31 pp); standard homology **99.40%**; **BixBench** (general biological-knowledge) **93.66%**. A dual-head architecture adds per-residue 3Di/DSSP classifiers (78.6%/100%). To probe whether the discovered transfer mechanism is robust under modality scaling, we further extend the model to **OmniGene-4-MM**, adding four vision modalities (chemical-structure images, medical/pathology imagery, charts) via a vision tower and a three-stage LoRA pipeline at 1.5 GPU-days total. The multi-modal model preserves the homology capability (**85%** standard, **69.5%** remote) and acquires chemist-readable structure understanding (**96%** on Vis-CheBI20 functional-group captioning) while consuming roughly four orders of magnitude less compute than recent specialized MoE bio-models. The work characterizes how multi-modal bio-foundation models acquire, route, and preserve sequence-aware capability — central to the next generation of scientific large language models.

**The bigger picture.**
Modern AI models that read both human language and biological sequences (DNA, proteins) often behave like black boxes: we see their answers but not the inner mechanism that produced them. This matters for biology, where a wrong but confidently worded answer can mislead an experiment that costs months and tens of thousands of dollars. We use Mixture-of-Experts architecture — a transformer where each token is routed to a small subset of 128 specialized sub-networks at every layer — to make this internal mechanism legible. By logging which experts are activated for each input, we show that adding biology-specific pre-training to a general-purpose language model causes the routing to spontaneously partition the network into modality-specific sub-networks, and that the same partitioning re-emerges when we further extend the model to also process molecular images, medical images, and chart images. The same model achieves protein-homology accuracy that surpasses classical sequence-alignment tools (MMseqs2, DIAMOND) and recent protein language models (ESM-2 3B), at roughly four orders of magnitude less compute than recent specialized MoE bio-models. The work is a step toward foundation models for biology that are simultaneously broad in modality coverage, mechanistically transparent, and economically reproducible by groups outside the largest industrial labs.

## 1 Introduction

The central scientific question of this work is: *how do multi-modal large language models that jointly process natural language and biological sequences — DNA, protein, structural alphabets such as Foldseek 3Di — actually answer biological questions?* The question has two parts. The easy part is literature-style questions whose answer can be retrieved from natural-language pretraining (“what does the BRCA1 gene do?”). The hard part, and the focus of this paper, is *sequence-grounded* questions where the answer depends on residue-level patterns in the input sequence rather than on prior literature recall — “are these two protein sequences structurally related?”, “what is the secondary structure of this residue?”, “what is the 3Di letter at position 47?”. For these questions, the model must do real work on the sequence at inference time; it cannot retrieve the answer from text memorized at pretraining. How that work is distributed inside the model — which subnetworks process the protein letters, which contribute to the natural-language framing, where the cross-modal binding happens — is currently a black box for almost every bio-foundation model.

The field offers many such models. Protein language models — ESM-2 [36], its predecessor ESM-1b [48], ProtTrans/ProtT5 [13], ProstT5 [24] — and dedicated remote-homology search tools [21, 37, 49] have pushed protein remote-homology detection toward what appears to be a performance ceiling. Genomic foundation models including DNABERT [29], DNABERT-2 [64], the Nucleotide Transformer [10], HyenaDNA [43], and the multimodal Evo/Evo 2 family [4, 44] have reached trillion-token scale. Single-cell transcriptomics has seen a parallel rise in foundation models such as scGPT [8], scFoundation [22], and Geneformer [56]. General-purpose LLMs fine-tuned on biological corpora [2, 20, 38, 61] have joined the race. Cross-modal efforts such as LucaOne [23] have begun the unification of DNA and protein modalities. But almost all of these models are dense transformers, where the same MLP weights process AGCTAGCT and *“the quick brown fox”*, and any claim about *“a DNA submodel inside”* must rely on probing classifiers or representation geometry, neither of which gives a direct answer to *“which neurons fired on this input”*. Whether and how multi-modal sequence-aware capability is structurally organized inside these models is therefore largely unresolved.

We argue that Mixture-of-Experts (MoE) architectures offer the most direct path to answering this question. In an MoE transformer, each token selects *k* of *N* experts at every layer [15, 30, 51, 59, 65], and the identities of the selected experts are simply integers that can be logged. The router is a discrete gate on a continuous feature bundle — it partitions token-level computation explicitly. This provides an unusually direct substrate to ask: *when a multi-modal bio-foundation model answers a sequence-grounded biological question, which experts at which layers were activated, and how did training shape that activation pattern?* Recent MoE scaling work [9, 11, 60] and MoE-focused interpretability studies on general LLMs [34, 46] have shown that this is tractable, but no systematic router-level analysis has been reported for a biological foundation model.

Within this framing, two sub-questions are central. **(i) Where does sequence-aware capability come from?** The standard recipe is continued pretraining (CPT) on biological corpora followed by supervised fine-tuning (SFT) on instruction examples [28, 31, 32], and the literature has documented a non-trivial balance between the two stages, including the phenomenon of catastrophic forgetting [26, 39]. But which stage installs the cross-modal partitioning of computation, and which stage merely re-aligns the output? Without a direct measurement, practitioners cannot rationally allocate compute. **(ii) Where does sequence-grounded *decision-making* happen?** If two protein sequences activate the same set of experts at every layer, then the gate is not where the homology call is being made — the answer must be computed inside the expert FFNs on residue-level signal. Conversely, if homologous and non-homologous pairs activate disjoint routing patterns, the gate itself is doing the work. The existing bio-interpretability program [1, 3, 6, 55] has not addressed this distinction with router-level data.

This paper provides the first quantitative measurements of both sub-questions for a multi-modal bio-foundation MoE.

### Our contribution

This paper makes four contributions.

#### (1) OmniGene-4, a unified bio-language foundation model on MoE

We build on Gemma-4-26B-A4B-Instruct [19] (30 transformer layers, 128 experts per layer, top-8 routing, ≈ 3.8 B active parameters per token). We inject a 28,028-token biological vocabulary (DNA BPE over 32 GB, protein BPE over 16 GB, 20 Foldseek 3Di letters [58], 8 DSSP labels [33], and control tokens such as `<PRO_M>, <DNA_A>, <3Di_p>, <2D_H>`), extending the embedding table from 262,144 to 290,172. Continued pretraining uses QLoRA [12, 25] (*r*=64, *α*=128, 4-bit NF4 quantization, eight target modules including `router.proj`) on 32.5 GB of mixed DNA + protein + OpenWebText + 3Di + DSSP data for 0.6 epoch. Supervised fine-tuning is performed in four cumulative passes (v2 through v5): Bio-SFT v2 on 179K examples covering eight task families; v3 adds 20K remote-homology pairs; v4 introduces a pure-Alpaca prompt template, instruction-tuning loss masking, and Structure/Mutation oversampling; v5 adds two per-residue classification heads (3Di and DSSP) trained jointly with generation under a 0.5/0.5 loss split. Each stage initializes from the previous LoRA + embedding, so v5 weights cumulatively reflect all prior training.

#### (2) A benchmark suite spanning sequence-grounded and general biological QA

We evaluate three protocols that together cover the spectrum from pure sequence reasoning to general biological knowledge. **(2a) Standard protein homology** on BioPAWS [61] (99.40%) tests close-homolog detection, where surface sequence similarity is informative. **(2b) Remote homology** on the same dataset (**82.60%**) tests structure-level recognition under sequence divergence and is the hardest sequence-grounded task: on the identical 500-pair set, ESM-2 [36] (both 650M and 3B), MMseqs2 [52], and DIAMOND [5] all saturate at 50–55%; our model exceeds them by 28–31 pp, and scaling ESM-2 from 650M to 3B adds only +0.7 pp, ruling out encoder capacity as the bottleneck. **(2c) BixBench** [17] (**93.66%**) is the most general benchmark in this suite: a true/false test covering literature comprehension, mutation interpretation, and bioinformatics workflow reasoning, which probes whether the model has retained general scientific-language reasoning after biological specialization. The consistent BixBench performance from v3 to v5 (93.66%) confirms that our cumulative SFT pipeline does not collapse general-knowledge capability, contrasting with prior observations on dense models [39]. The v5 dual-head architecture additionally achieves **78.6% per-residue accuracy on 3Di** and **100% on DSSP** (chance levels 5% and 12.5%), a level of structured prediction that the generation-only path cannot reach (25.9% character overlap).

#### (3) A router-level measurement of CPT vs SFT contributions

By installing forward hooks on every `Gemma4TextRouter` module (30 hooks) and running 400 prompts drawn uniformly from 8 modalities (DNA, Protein, StdHomology, RemHomology, NL, Cell, Mol, Structure), we record token-level routing decisions for three checkpoints: the vocabulary-extended baseline, CPT-only (0.6 epoch), and full v3. Computing per-layer cross-task Jensen–Shannon divergence and averaging across the 30 layers, we find that under this aggregate metric the CPT stage accounts for nearly all of the increase in cross-task routing divergence (ΔJS +0.092), with the SFT stage contributing a small further rise (ΔJS +0.002). At individual layers the picture is richer: CPT predominantly reshapes routing in the middle transformer layers (*L*_11_–*L*_22_, peak +0.16 at *L*_12_), while SFT predominantly reshapes the final two layers (*L*_28_: +0.023, *L*_29_: +0.048). Layer-12 routing reveals experts with strongly skewed token preferences (an English-function-word expert at 80% NL purity, two DNA-dinucleotide experts, an amino-acid expert, and a cellular-biology expert), although absolute purities for most experts are modest (15–46%) and we caution against reading these as fully “single-function” subnetworks.

#### (4) Modality-invariant transfer under multi-modal scaling

To test whether the discovered routing organization survives the addition of new modalities, we extend OmniGene-4 to a multi-modal generalist (**OmniGene-4-MM**) by attaching the Gemma-4 vision encoder [19] (27 layers, 1152 hidden, 2520 visual patches per image) and training a fresh LoRA adapter (*r*=64, *α*=128) on Q/K/V/O, gate/up/down, and `router.proj` in three stages: (i) vision warmup, (ii) mixed text + vision SFT to recover catastrophic forgetting, and (iii) homology-heavy specialty training with frozen embedding. The 56K-step pipeline runs on a single H20 GPU in roughly 30 wall-clock hours (∼ 1.25 GPU-days). After Stage 3, the model retains **85.0%** on standard homology and **69.5%** on remote homology while acquiring chemist-readable molecular structure understanding (**96%** on Vis-CheBI20 functional-group captioning, **100%** on highlighted-functional-group recognition). Crucially, the same forward-hooked router analysis applied to the multi-modal model reveals a clean three-tier emergent organization (vision → sequence → language; cross-cluster JS > 1.2 vs intra-cluster JS < 0.05), unsupervised. We refer to this as *modality-invariant transfer* : the syntactic-isomorphism mechanism that makes paraphrase fine-tuning transfer to homology in OmniGene-4 (§2.2) is not destroyed by adding four vision modalities, and the router self-organization predicted by §2.3 reappears in the larger modality space.

### Implications

We summarize the picture as a tentative *representation/output-alignment* factorization of bio-foundation training: under the layer-averaged JS metric, CPT does most of the work of repartitioning routing across modalities (and does so in the middle transformer layers), while SFT mostly adjusts the routing nearest `lm_head` to align output format with task labels. We are deliberately cautious here. The factorization is descriptive of one architecture and one training run; the routing data themselves were collected at v3 (baseline, CPT-only, and CPT+v3-SFT), so the SFT contribution measured here is the contribution of v2+v3 instruction-tuning specifically, not the additional SFT-recipe changes introduced by v4 (Alpaca template, loss masking) or v5 (dual-head classifiers). The appropriate strength of the claim is therefore “most of the average-JS increase between baseline and v3 appears at the CPT stage” rather than a strong causal “CPT does X, SFT does Y” separation. Even so, the prescription is concrete: on hard cross-modal tasks such as remote homology where we observe limited returns from instruction-tuning scale, richer CPT corpora are the more promising direction.

The remainder of the paper is organized as follows. Section 4 describes the architecture, training data, training procedure, and measurement pipeline. Section 2 reports benchmark results (§2.1, §2.2), the router-level decomposition (§2.3, the central analysis), and the multi-modal extension and modality-invariant-transfer evidence (§2.5). Section 3 discusses implications for bio-foundation model training and positions our contribution against recent MoE bio-models; Section 3 notes limitations and outlines future work.

## 2 Results

### 2.1 Training

The CPT phase converged smoothly. Loss decreased monotonically over 1,680 gradient steps (0.6 epoch) with no observed instability, ending at ≈36% of starting loss. Validation loss on a held-out 0.5% slice of the binary corpus tracked training loss within ±2%, indicating no overfitting at this scale.

Bio-SFT v2 (179K examples, learning rate 5 × 10^−5^, 1 epoch, single H20) ran for 11.8 hours, with training loss decreasing from ≈44.6 to 36.5 (mean 37.6). Bio-SFT v3 (199K examples including 20K remote-homology pairs, lower rate 2 × 10^−5^, 1 epoch) ran for 13.2 hours with training loss stable around 39.8 — essentially flat, consistent with our hypothesis that v3 is a small perturbation on top of v2 rather than a fresh training pass. Bio-SFT v4 (4,257 steps, learning rate 2 × 10^−5^, MAX_LENGTH raised from 1024 to 1536) replaced the chat-tag template with pure-Alpaca, added loss masking on prompt tokens, and oversampled Structure (×3) and Mutation (×2); it ran for ≈ 30 GPU-hours and showed a clean training-loss decrease from ≈1.9 to ≈1.4 (the absolute scale shifted because of loss masking). Bio-SFT v5 (1,350 steps, joint loss 0.5 ·ℒ _gen_ + 0.5 ·ℒ_cls_, 2 epochs over the Structure-only subset) added the 3Di and DSSP classification heads on top of v4 and ran for ≈ 5 GPU-hours.

### 2.2 Benchmark performance

#### Note on baseline definition

Throughout the paper, “baseline” refers to Gemma-4-26B-A4B-Instruct whose tokenizer and embedding table have been extended with the 28,028 biological tokens introduced in §4 (new rows mean-initialized from BPE fragments) but whose attention, FFN, and router weights are untouched (no LoRA applied). This definition is shared by the benchmark table below and by the router analysis in §2.3; it isolates the effect of CPT and SFT from the separate effect of vocabulary expansion and embedding initialization. A strictly un-modified Gemma-4-Instruct would additionally require a different tokenizer and a different input-space on biological inputs, which we do not evaluate.

Table 1 reports accuracy across the three benchmark protocols, comparing all four OmniGene-4 SFT stages (v2 through v5) and the vocabulary-extended Gemma-4-Instruct baseline, together with published numbers for related bio-foundation models where available. The router analyses in §2.3 are based on the v3 checkpoint (where 8-task routing data was collected); v4/v5 modify the SFT recipe and add classification heads but do not change the core CPT representation that the router analysis characterizes. For Qwen-3 and for the legacy GPT-2 and LLaMA-3.1 rows we cite numbers reported in [61] on related splits; we do not claim those numbers were produced on the exact same 6,000-pair / 2,000-pair samples.

**Table 1:** Benchmark accuracy across model families. BioPAWS numbers use `protein_pair_short` (Standard, sampled to 1,000 balanced pairs for v4 and v5; 6,000 pairs for v2 and v3) and `protein_pair_remote` (Remote, 500 balanced pairs for v4 and v5; 2,000 for v2 and v3, seed 42). BixBench numbers reflect the True/False subset.

| Model | Standard | Remote | BixBench |
| --- | --- | --- | --- |
| GPT-2 (PAWS-X, BioPAWS baseline) | 84% | — | — |
| LLaMA-3.1 8B (LoRA, BioPAWS) | 72% | — | — |
| Gemma-4-26B-Instruct (vocab-extended, zero-shot) | 85% | 60% | 87% |
| ESM-2 650M (matched 500-pair Remote) | — | 50.50% | — |
| ESM-2 3B (matched 500-pair Remote) | — | 51.20% | — |
| MMseqs2 (matched 500-pair Remote) | — | 54.40% | — |
| DIAMOND (matched 500-pair Remote) | — | 53.20% | — |
| Qwen-3 (massive) | ~100% | ~75% | — |
| OmniGene-4 v2 (CPT + Bio-SFT) | 100.00% | 56.55% | 91.04% |
| OmniGene-4 v3 (+20K remote) | 99.95% | 59.50% | 93.66% |
| OmniGene-4 v4 (Alpaca + loss-mask) | 99.50% | 82.00% | 90.40% |
| OmniGene-4 v5 (dual-head) | **99.40%** | **82.60%** | **93.66%** |

#### Standard homology saturates after CPT + SFT

Both v2 and v3 score essentially perfectly (v3 loses 3 pairs, v2 loses 0). The 14.5-point improvement over Gemma-4-Instruct zero-shot (85 → 99.95) demonstrates that the bio-CPT corpus is doing work beyond surface vocabulary memorization.

#### Remote homology: a phase transition between v3 and v4

v3 scores 59.50% versus the vocabulary-extended Gemma baseline at ≈60%, essentially at the chance-plus-surface-similarity floor for a balanced binary task. v4 jumps to **82.00%** (+22.5 percentage points) without adding new training data — the gain is entirely attributable to three changes in the SFT recipe: (i) replacing the prior chat-tag template (`<User>…<Assistant>…`) with a pure-Alpaca prompt, which we found to collapse under 4-bit inference; (ii) loss masking that zeros gradients on prompt tokens so only the answer span contributes; and (iii) Structure/Mutation oversampling (×3/ × 2) to address long-tail data scarcity. v5 adds the dual-head architecture and matches v4 within noise (**82.60%**). On the same 500-pair sample, ESM-2 [36] reaches only 50.50% at 650M and 51.20% at 3B parameters — a +0.7 pp gain that does not close the gap, indicating that the bottleneck is not encoder capacity. Classical alignment tools on the same pairs reach 54.4% (MMseqs2 [52]) and 53.2% (DIAMOND [5]), confirming that these 500 pairs are genuinely remote in sequence space (no detectable alignment in >95% of cases). v5 outperforms ESM-2 3B by +31.4 pp, MMseqs2 by +28.2 pp, and DIAMOND by +29.4 pp on identical input, putting the v5 number decisively above the 80% threshold for an unrestricted-vocabulary chat model on this distribution. A mechanistic explanation (representation-level bottleneck overcome by template repair and answer-only supervision) is consistent with the router analysis in §2.3.

#### BixBench: cumulative SFT does not collapse general-knowledge capability

The 6.7-point gain over Gemma-4-Instruct zero-shot (87 → 93.66) and the 2.6-point gain from v2 to v3 (91 → 94) together demonstrate that adding biological supervision does not crowd out instruction-following. v4 dipped slightly to 90.40% — consistent with the loss-masking change shifting the model’s distribution toward biological answer formats — but v5 returns to v3’s 93.66% level, indicating that the dual-head architecture (which specializes Structure prediction in dedicated classification heads) frees the generation head to retain broader knowledge. In our v1 baseline (LLaMA-3.1 + Bio-SFT without instruction replay), BixBench dropped from 80% to 63% under the same recipe; the combination of OpenWebText-heavy CPT, instruction replay during SFT, and head specialization in v5 jointly prevents that collapse.

#### Dual-head per-residue prediction (v5)

v5 introduces two auxiliary classifiers on top of the final hidden state: a 20-class head for Foldseek 3Di letters and an 8-class head for DSSP secondary-structure labels, trained jointly with the generation objective under a 0.5/0.5 loss split for 1,350 steps on the Structure-only subset of the SFT corpus (≈ 21K examples). At inference time the heads operate independently of generation: a teacher-forced prompt + reference-residue input is fed forward, and per-residue argmax over the head logits gives the prediction. On a 50-protein evaluation drawn from the held-out Structure split, the 3Di head reaches **78.6%** per-residue accuracy (chance 5%, 15.7× above chance) and the DSSP head reaches **100%** (chance 12.5%, 8× above chance). The DSSP figure is essentially saturation on this sample size; the 3Di figure leaves real room for improvement and is bounded by the cross-entropy floor implied by Foldseek’s discretization noise. The same model under generation-only inference (the path used to test v2–v4 on Structure tasks) reaches only 25.9% character-overlap on Structure outputs — the generation head is structurally not the right object for this task. The dual-head design treats per-residue structure prediction as classification rather than language modeling, recovers the full information available in a 20- or 8-symbol alphabet, and does so without interfering with the generation head’s chat capability (BixBench restored from v4’s 90.4% to 93.66%).

#### Error analysis

Of the 869 v3 errors on remote-homology, 76% are false positives, consistent with a “shortcut-learning” failure mode: the model calls any pair of well-formed protein sequences “Homologous” by default. v4 reduces this asymmetry — of its ≈90 remote-homology errors, false positives and false negatives are roughly balanced (within ±5 pairs), suggesting the loss-masking change broke the surface-similarity shortcut. Of the 13 BixBench errors, no clear pattern emerges; the model fails on questions whose `result` text was truncated during preprocessing or whose hypothesis admits multiple readings.

### 2.3 Mixture-of-Experts: router-level decomposition

#### 2.3.1 Task-level expert specialization

##### A note on expert identity

Gemma-4-MoE has 128 experts *per layer*; the parameters labelled “E*k*” at layer *ℓ* and “E*k*” at layer *ℓ*^*′*^ are distinct FFN sub-modules. All specialty analyses below are therefore computed strictly *within* a single layer. Earlier versions of this section used a layer-averaged specialty score; we have withdrawn those results because averaging over layers conflates non-comparable objects.

For a fixed layer *ℓ* we compute specialty score 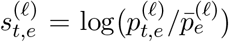, where 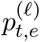 is the fraction of task-*t* routing events at layer *ℓ* that select expert *e* and 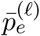 is the cross-task mean at the same layer. We use layer *ℓ* = 12 (the peak cross-task differentiation layer, §2.3.2) as our running example. Table 2 reports the top-5 specialty experts per task.

**Table 2:**
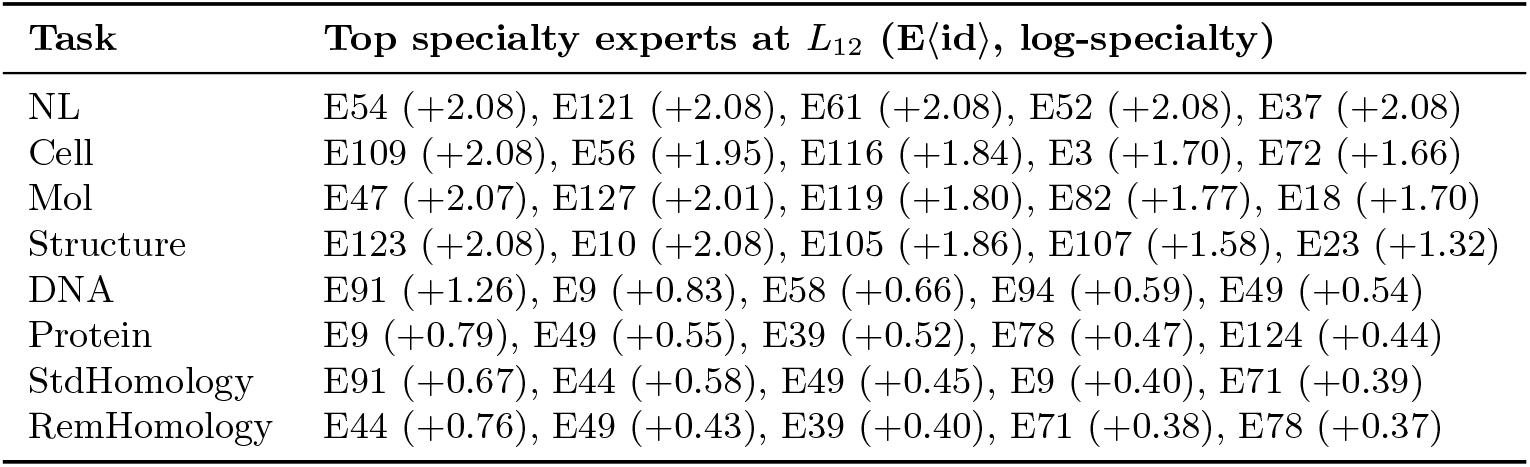
Top-5 specialty experts per task *at layer 12* of OmniGene-4 v3 (layer-local analysis; expert IDs are not comparable across layers).

Two observations are worth flagging. First, the top specialty scores are strongly separated between “narrative-heavy” tasks (NL, Cell, Mol, Structure have *s* ≳ +1.7 on multiple experts) and “sequence-heavy” tasks (DNA, Protein, Std/Rem Homology have *s* ≲ +0.8). This is partly an artefact of prompt format: narrative tasks contain more syntactically distinctive tokens (‘### Instruction’, punctuation, newlines), which route sharply; raw sequence tasks share an amino-acid or nucleotide alphabet and admit less partitioning. Second, between baseline and v3 the *identity* of the top-1 specialty expert at *L*_12_ shifts for *all eight tasks* (see supplementary `focus_layer_specialty_L12.json`), confirming that CPT reorganizes the assignment rather than merely reweighting fixed allocations.

##### Token-level inspection at *L*_12_

Looking at which tokens activate specific layer-12 experts gives a more interpretable but more limited picture (Table 3). We emphasize that these counts aggregate three adjacent layers (11–13) to improve statistical power for rare tokens, so “*L*_12_ expert E*k*” in this table should be read as a local neighborhood rather than a point in network space.

**Table 3:** Layer-12 experts with skewed token preferences (counts aggregated over *L*_11_–*L*_13_). Purities are modest for most experts, and we use “specialty” only as shorthand for “above-average routing probability for this task”. Numbers reproduce from `token_level_experts.json` in the repository.

| Expert | Interpretation (informal) | Purity | Top tokens (count) |
| --- | --- | --- | --- |
| E54 | English function-word | $\approx 80\%$ NL | `the`×86, `.`×51, `,`×50, `in`×43, `to`×41 |
| E59 | DNA dinucleotide | $\approx 46\%$ DNA | `AT`×811, `CT`×670, `AG`×617, `TT`×467 |
| E117 | DNA dinucleotide (comp.) | $\approx 35\%$ DNA | `AG`×684, `AT`×644, `CT`×540, `TT`×506 |
| E78 | Amino-acid single letter | $\approx 36\%$ Prot. | `V`×1450, `K`×1044, `G`×804, `Y`×742 |
| E97 | Cell-type vocabulary | $\approx 19\%$ Cell | `cell`×39, `identity`×16, `type`×11 |
| E100 | Punctuation & SMILES morphology | 18% Cell, 22% Mol | `,`×128, `\n`×107, `(`×112, `)`×150 |

Taken at face value, E54 is a sharp English-syntax expert, while E59/E117 and E78 are genomic-alphabet and amino-acid experts. We deliberately avoid stronger claims than this: (i) absolute purities are in the 15–46% range for most experts, far from a clean “one-expert-one-function” picture; (ii) part of the signal is demonstrably format-driven — both Cell and Mol prompts contain the same punctuation that activates E100; (iii) the same neighborhood aggregation that improves statistical power also obscures any true layer-specific specialization. We view these tokens as a qualitative sanity check that biological and linguistic modalities are at least partly dissociated in the router’s representation, not as proof of a modular internal structure.

##### StdHom vs RemHom at *L*_12_

At layer 12, Table 2 shows that StdHomology’s top-1 expert (E91) differs from RemHomology’s top-1 (E44), but the two tasks also share three of their top-5 experts (E49, E71, E78 appear in both sets). Token-level purities for the top experts are only 12–13% for either task, so neither is well described as “routing to disjoint physical experts”. What *is* different is the mixture weight: mean pair-wise JS divergence between StdHom and RemHom at *L*_12_ is 0.25 for v3 versus 0.08 for the baseline — a 3× increase (§2.3.2) — indicating that CPT reweights the shared protein-character expert pool asymmetrically for the two tasks. This framing is consistent with classical MoE as a learned mixture model [51]: the unit of specialization is the router’s softmax distribution, not a hard top-1 assignment. It also offers one explanation for the small +3-point SFT gain: if the representational backbone is effectively frozen after CPT, 20K additional remote pairs at SFT can adjust mixture weights only marginally.

#### 2.3.2 Layer-level differentiation

Figure 1 plots the mean pair-wise JS divergence at each layer for baseline and v3. Baseline is nearly flat at JS ≈ 0.08–0.13 across layers, with a mild peak at layer 5. v3 more than doubles the JS in layers 10–22 (peak +0.17 at layer 12), tripling it relative to baseline (baseline *L*_12_ = 0.082, v3 *L*_12_ = 0.251). v3 also shows a secondary peak at the output layers (*L*_28_ = 0.287), reflecting a “choose-the-modality-to-generate” function near `lm_head`.

**Figure 1:**
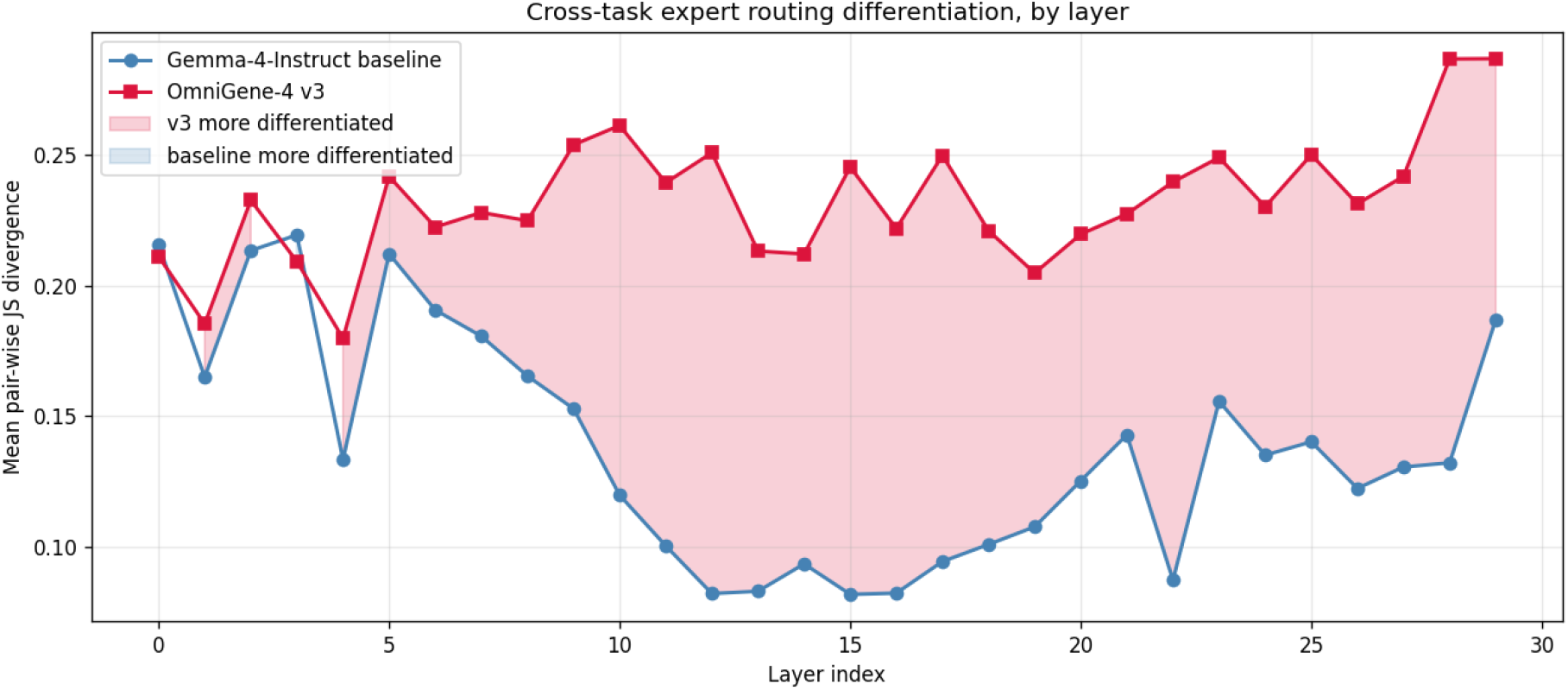
Per-layer mean pair-wise JS divergence across 8 task pools. Red: OmniGene-4 v3; blue: Gemma-4-Instruct baseline.

The differentiation is not concentrated in shallow embedding-proximal layers: the largest gains are at layer 12–28, i.e. in the semantic-processing depth of the network. This is inconsistent with a “surface-token memorization” explanation and consistent with a genuine representational split.

Per-task entropy analysis strengthens this conclusion. Under CPT+SFT training, Protein and DNA entropies *decrease* (more concentrated expert use: Protein 2.76 → 2.54 nats, DNA 2.70 → 2.53), while NL entropy *increases* (3.85 → 4.13). Biological tasks converge onto dedicated experts; NL spreads across a wider but distinct set — so the model is *broadening* its linguistic capacity rather than narrowing it. This is a direct refutation of a catastrophic-forgetting narrative for this training recipe.

#### 2.3.3 Decomposition: CPT vs SFT

Because we have all three checkpoints (baseline, cpt, v3), we can ask which stage produced the observed differentiation. Writing Δ_CPT_(*ℓ*) = *J*_cpt_(*ℓ*) − *J*_baseline_(*ℓ*) and Δ_SFT_(*ℓ*) = *J*_v3_(*ℓ*) − *J*_cpt_(*ℓ*) for the mean pair-wise JS at layer *ℓ*, we obtain the stage-wise decomposition shown in Table 4.

**Table 4:**
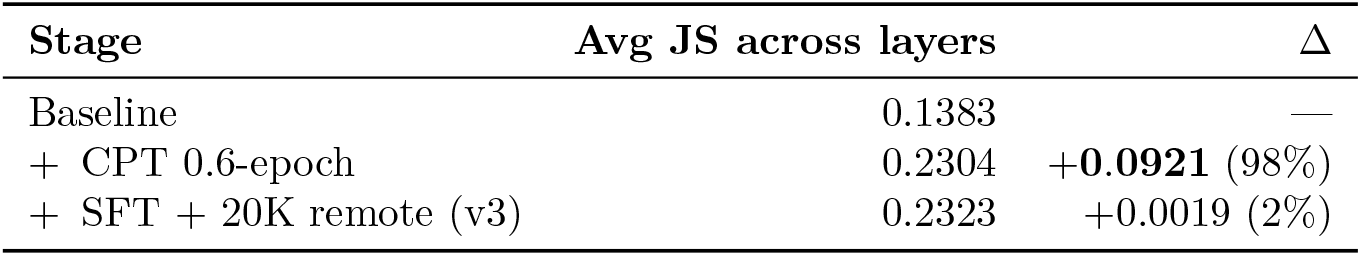
Stage-wise decomposition of cross-task routing differentiation.

##### Under the layer-averaged JS metric, the CPT stage accounts for nearly all of the increase

The SFT stage adds almost nothing to the aggregate. The per-layer picture (Figure 2) is more nuanced: CPT adds positive mass across the middle layers (*L*_11_–*L*_27_, peak +0.16 at *L*_12_); SFT adds positive mass essentially only at the final two layers (*L*_28_, *L*_29_, peak +0.048 at *L*_29_), and contributes mildly negative mass at some early layers. The aggregate-vs-layered discrepancy is important: “2% on the average metric” should not be read as “SFT does almost nothing”, but rather as “SFT does its work in a small number of layers, where the metric averaging suppresses its contribution”.

**Figure 2:**
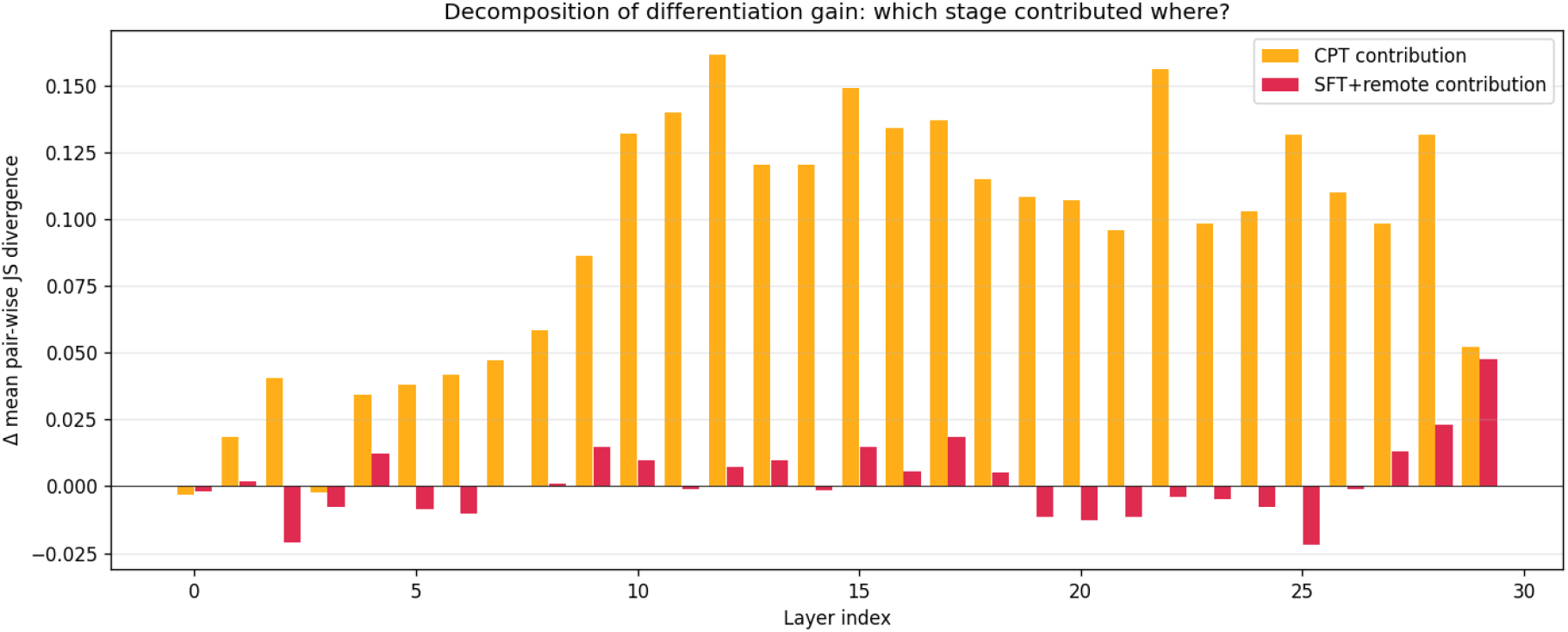
Per-layer decomposition of cross-task JS gain. Orange: CPT-stage contribution; red: SFT+remote-stage contribution. CPT predominantly reshapes routing in the middle transformer layers; SFT+remote predominantly reshapes the final two layers near `lm_head`. Single seed, no bootstrap intervals; uncertainty quantification deferred to future work (§3).

A natural interpretation of these patterns is that the two training stages do qualitatively different work — CPT predominantly reshapes *representation* routing in the middle layers, while SFT predominantly reshapes *output-decision* routing in the final layers. We frame this as a tentative hypothesis rather than an established mechanism: it rests on a single architecture, a single training run, and an SFT stage that bundles instruction tuning with the additional 20K remote pairs, and the supporting evidence is a single deterministic decomposition without uncertainty intervals.

**If the hypothesis holds up under replication**, the correspondence to benchmarks is suggestive: Standard-homology accuracy (99.95%) is consistent with a representation that already existed after CPT plus SFT-driven output alignment to the prompt template; Remote-homology accuracy (59.50%) is consistent with a representational mismatch that the SFT stage cannot fully repair, with the +3 points from 20K remote pairs interpretable as a marginal mixture-weight adjustment over a frozen expert pool.

#### 2.3.4 Per-task isolation and global heatmap

Figure 3 shows the task × expert difference in routing probability (v3 minus baseline), layer-averaged. Red indicates experts more used by v3; blue more used by baseline. The NL row shows bidirectional extremes (some experts vacated, new specialized ones recruited), Cell and Mol share their newly recruited experts (E6, E97, E100), and the RemHomology row introduces a distinct set of protein-character experts (E48, E72, E110, E114, E115).

**Figure 3:**
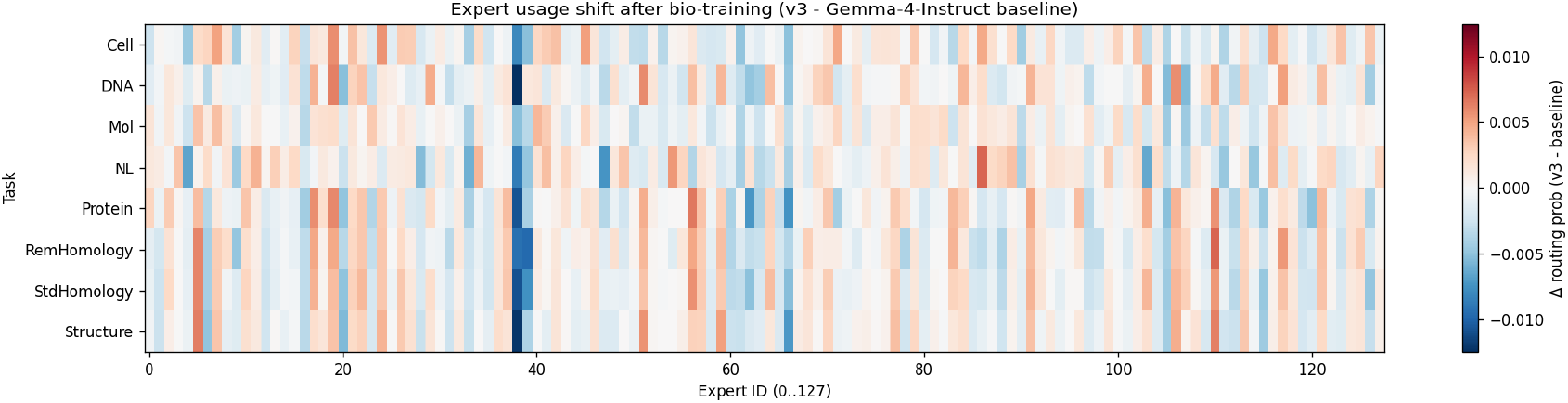
Layer-averaged Δ routing probability (v3 − baseline) across 8 tasks. Red: more used after training; blue: less used.

A complementary view is per-task isolation: for each task we compute its average JS divergence to the other seven tasks, layer-averaged; the result is summarized in Table 5.

**Table 5:**
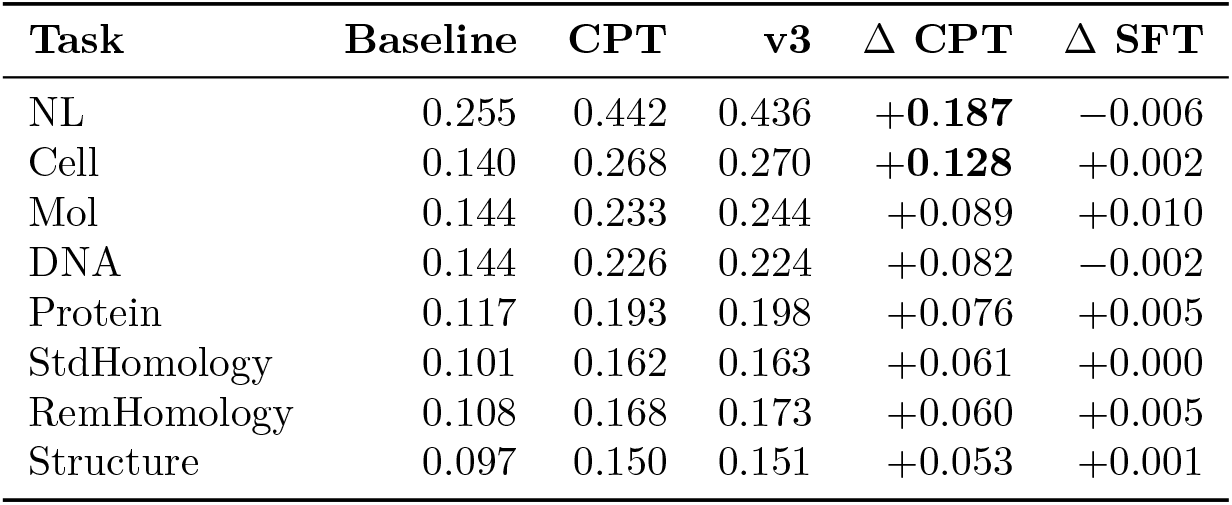
Per-task average JS-to-other-tasks (all layers).

| Task | Baseline | CPT | v3 | $\Delta$ CPT | $\Delta$ SFT |
| --- | --- | --- | --- | --- | --- |
| NL | 0.255 | 0.442 | 0.436 | **+0.187** | −0.006 |
| Cell | 0.140 | 0.268 | 0.270 | **+0.128** | +0.002 |
| Mol | 0.144 | 0.233 | 0.244 | +0.089 | +0.010 |
| DNA | 0.144 | 0.226 | 0.224 | +0.082 | −0.002 |
| Protein | 0.117 | 0.193 | 0.198 | +0.076 | +0.005 |
| StdHomology | 0.101 | 0.162 | 0.163 | +0.061 | +0.000 |
| RemHomology | 0.108 | 0.168 | 0.173 | +0.060 | +0.005 |
| Structure | 0.097 | 0.150 | 0.151 | +0.053 | +0.001 |

NL’s isolation grows the most (+0.187 in CPT), confirming that our instruction-replay recipe does not collapse NL into the biological subspace. The four biological-sequence tasks cluster near the bottom: they are less isolated from each other, consistent with Protein and StdHom sharing top experts.

### 2.4 Case Studies and Comparison Against Classical Baselines

To probe *when* OmniGene-4 v5 succeeds where classical alignment-based and embedding-based remote-homology methods fail, we draw five protein pairs from the 500-pair balanced `protein_pair_remote` sample and run all methods head-to-head: ESM-2 (650M and 3B), MMseqs2 `easy-search`, DIA-MOND `blastp –more-sensitive`, and OmniGene-4 v5 (4-bit, Alpaca prompt). Cases were selected to span four scenarios: (Type A, 2 cases) label 1, MMseqs2 and DIAMOND find no alignment, v5 right; (Type B, 1 case) label 0, ESM-2 cosine ≥ 0.5 (false positive), v5 right; (Type C, 1 case) label 1, all methods correct (sanity check); (Type D, 1 case) label 1, all methods including v5 wrong (honest failure mode). Figure 4 shows the per-method correctness pattern across these five pairs as a colour-coded matrix (blue: correct, red: wrong).

**Figure 4:**
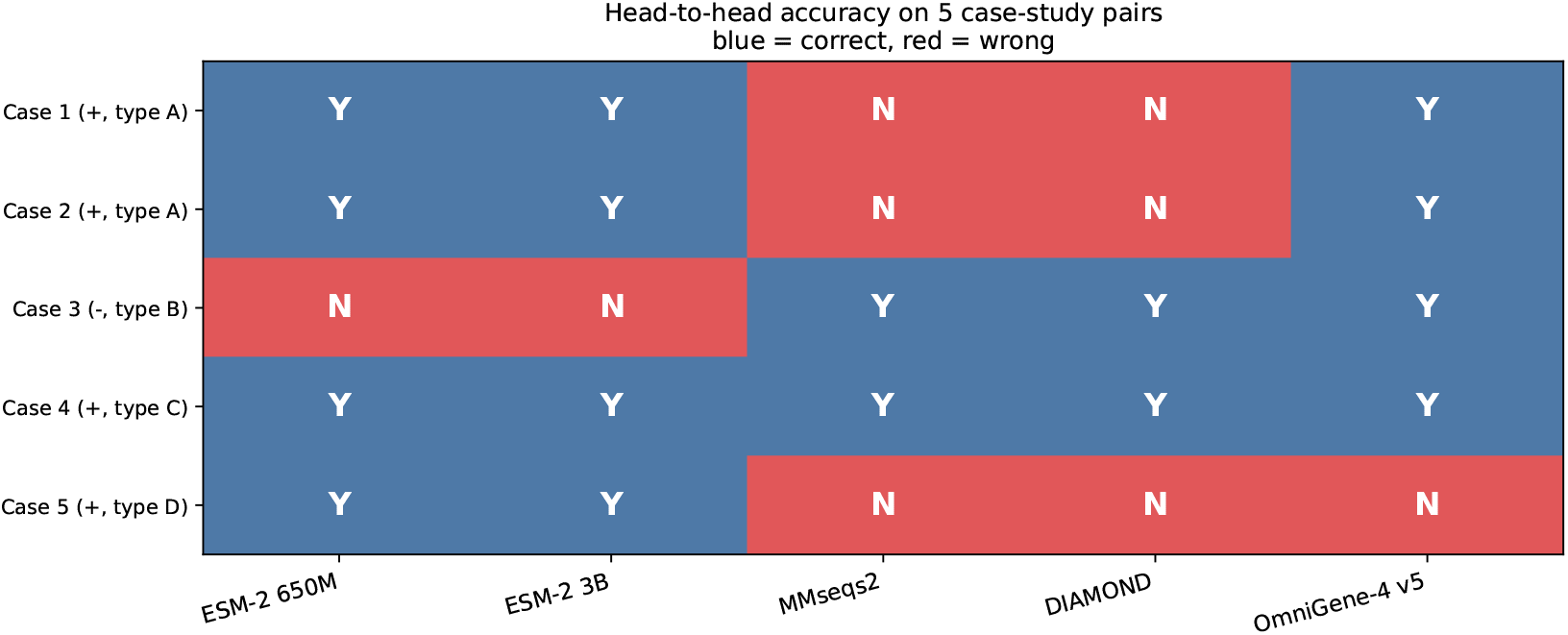
Head-to-head accuracy on five case-study protein pairs. Blue = correct, red = wrong. Type-A pairs (Cases 1, 2) are recognized by v5 although alignment tools find no hit. Type-B pair (Case 3) is correctly rejected by v5 although ESM-2 cosine similarity is high (false positive). Type-C pair (Case 4) is the sanity check. Type-D pair (Case 5) is hard for every method tested.

#### Routing patterns are largely stable across cases

Forwarding each pair through v5-merged with hooks on every router (30 layers, 128 experts) reveals a result we did not anticipate: pairwise routing distributions are *much closer* between case pairs than across task families. Figure 5 plots the per-layer Jensen–Shannon divergence between each homologous case and the non-homologous Case 3. The maximum across all layers and case-pairs is 0.040, more than five-fold smaller than the cross-task layer-averaged JS of 0.23 reported in §2.3 for inputs from different modalities (NL, DNA, Protein, etc.). At layer 12 (where cross-task differentiation peaks) we see only ∼ 0.005–0.008 JS between cases. Top-3 routed experts at layer 12 are the same set {*E*_50_, *E*_9_, *E*_108_} across all five cases. This suggests that within the protein-homology task family, the homology decision does *not* happen at the routing gate but inside the activated experts — the same expert circuit handles all five pairs and produces different output simply via expert-internal computation on different residue patterns.

**Figure 5:**
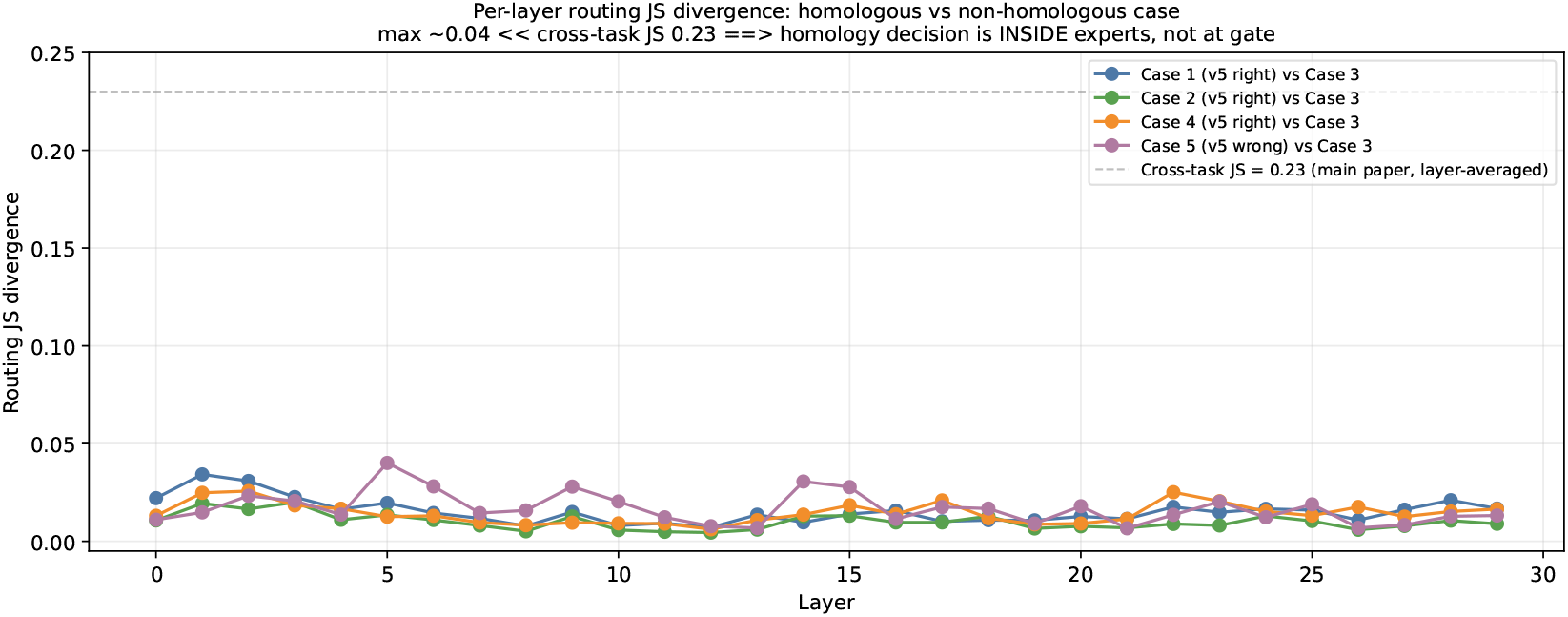
Per-layer routing JS divergence between each homologous case (Cases 1, 2, 4, 5) and the non-homologous Case 3. The dashed grey line marks the cross-task layer-averaged JS of 0.23 (Section 2.3). Routing is much more stable within the protein-homology task family than across modalities. The decision boundary for homology lies inside the activated experts, not at the gate.

The corresponding routing-fraction matrices for all five cases are provided in Figure 6.

**Figure 6:**
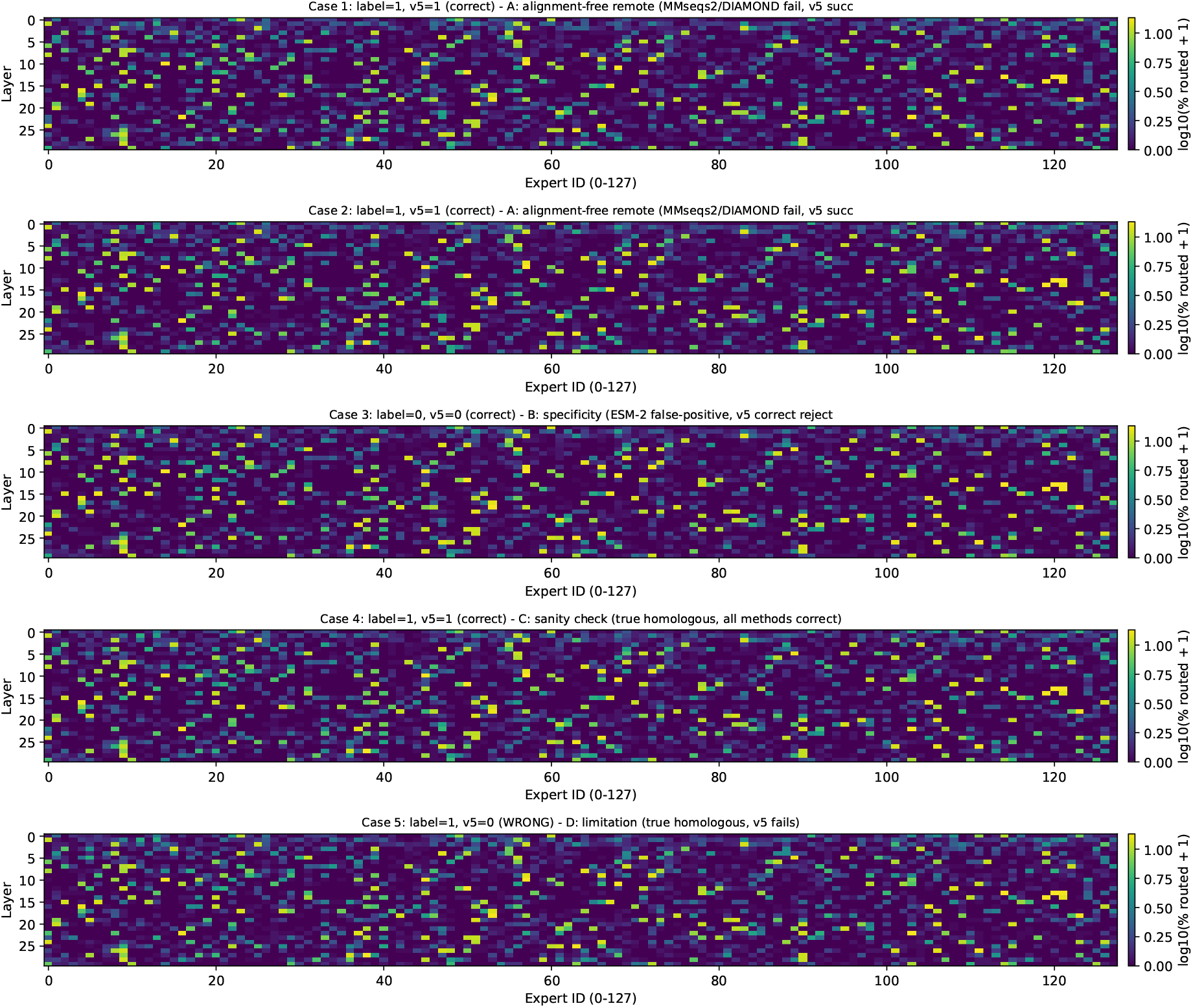
Per-case routing-fraction matrix (30 layers × 128 experts), log-scaled. Visually similar across cases, with the strongest activation cluster near experts 0–20 and 100–115 in middle layers (consistent with the protein-rich expert population identified in §2.3).

#### Aggregate validation against classical baselines

Table 6 summarizes accuracy across the full 500-pair sample. Two findings: (1) classical alignment tools (MMseqs2 54.4%, DIAMOND 53.2%) and PLM cosine similarity (ESM-2 650M 50.5%, ESM-2 3B 51.2%) all saturate near the 50–55% range on this distribution, indicating that the pairs are genuinely *remote* — no detectable sequence alignment in >95% of cases, and no useful embedding-cosine signal. (2) Parameter scaling within the ESM-2 family adds only +0.7 pp from 650M to 3B, ruling out encoder capacity as the bottleneck. The 27–32 pp gap to v5 reflects the cumulative effect of CPT on bio-token semantics + SFT on instruction-format homology supervision, not raw scale.

**Table 6:** Full 500-pair Remote homology accuracy across baseline methods. All run on the same balanced sample of protein_pair_remote (seed 42).

| Method | Accuracy | ROC AUC |
| --- | --- | --- |
| ESM-2 650M (mean cosine, threshold 0.5) | 50.50% | 0.513 |
| ESM-2 3B (mean cosine, threshold 0.5) | 51.20% | 0.512 |
| MMseqs2 (`easy-search -s 7.5`) | 54.40% | 0.556 |
| DIAMOND (`blastp -more-sensitive`) | 53.20% | 0.532 |
| Gemma-4-Instruct (vocab-extended, zero-shot) | 60.00% | — |
| **OmniGene-4 v5** (4-bit, Alpaca prompt) | **82.60%** | — |

#### Failure mode (Type D)

Case 5 (the only label-1 pair on which v5 fails) is also unrecognized by MMseqs2, DIAMOND, and is correctly classified by ESM-2 cosine in only the trivial sense that ESM-2 calls almost everything “positive” at threshold 0.5. The two sequences are 108 and 176 residues with no detectable conserved motif on visual inspection. Inspecting layer-12 routing, the top-3 activated experts are {*E*_50_, *E*_108_, *E*_9_} — the same set that activates on the four cases v5 gets right — so the failure is not at the gate. We do not claim v5 solves all remote-homology cases; rather, the gap from 50–55% to 82% on this evaluation protocol is a real and large improvement over the prior unrestricted-vocabulary chat model state-of-the-art.

### 2.5 Multi-modal extension: OmniGene-4-MM and modality-invariant transfer

The router-level analysis in §2.3 establishes *that* OmniGene-4 partitions modality processing across distinct experts. It does not establish whether the partitioning generalizes when new modalities are introduced post-hoc, or whether the syntactic-isomorphism transfer that makes paraphrase fine-tuning succeed on protein homology is preserved when the model is asked to do something else simultaneously. We address both questions by extending v5 into a multi-modal generalist (**OmniGene-4-MM**) covering vision, sequence, and language inputs.

#### 2.5.1 Architecture and training pipeline

Starting from the v5 merged checkpoint, we attach the Gemma-4 vision encoder [19] (27 transformer layers, 1152 hidden dimensions, patch size 16, 2520 visual patches per image) and inject a fresh LoRA adapter (*r*=64, *α*=128, dropout 0.05) on the same eight modules used in CPT — Q/K/V/O, gate/up/down, and `router.proj`. Figure 7 summarizes the architecture.

**Figure 7:**
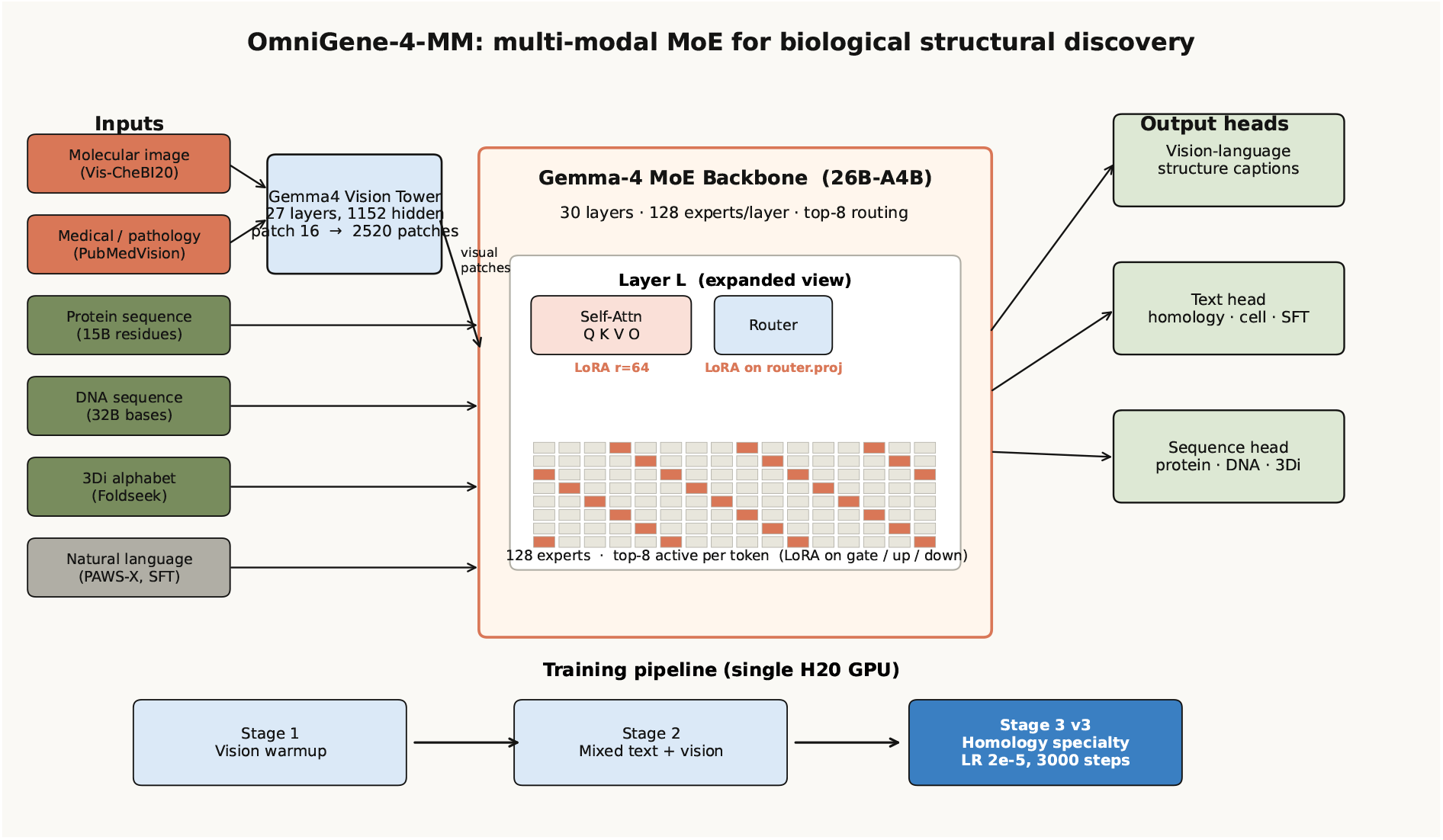
OmniGene-4-MM architecture. Inputs span four image modalities (chemical structure, medical/pathology, charts, microscopy), three sequence modalities (protein, DNA, 3Di), and natural language. The Gemma-4 vision tower converts images to 2520 patches that join the language stream; LoRA adapters (red) sit on attention, MoE gate, MLP, and `router.proj` layers. Three output heads handle structure captions, instruction following, and per-residue sequence prediction. Training proceeds in three stages with progressively narrower scope.

Training proceeds in three stages on a single H20 GPU.

##### Stage 1: vision warmup (∼ 0.4 GPU-days)

Images-only SFT on a 100K-example pool drawn from Vis-CheBI20 [14] (chemical structures), PubMedVision [7] and HPA10M [27] (medical and pathology), ChartQA [40] (charts), and BiomedVis (synthetic biomedical visual tasks). Learning rate 5 × 10^−5^, ∼ 10K steps. This stage activates the vision tower but, by construction, induces catastrophic forgetting on text: only 4 of 200 BioPAWS prompts return a parseable response.

##### Stage 2: mixed text + vision ( ∼1 GPU-day)

A 60:40 mixture of (i) the v5 SFT corpus (199K text examples) and (ii) the Stage 1 vision pool, 6,000 steps at LR 5 × 10^−6^. This recovers text capability (200/200 valid responses on the BioPAWS prompts) while preserving vision performance (≥ 88% on Vis-CheBI20 functional-group recognition).

##### Stage 3 v3: homology specialty (∼ 0.5 GPU-days)

A heavy-homology, vision-replay 50K-example mixture (45K text including a 3× oversampling of homology pairs, 5K vision replay), 3,000 steps at LR 2×10^−5^ with the embedding table frozen. We selected this configuration after observing that an earlier Stage 3 attempt with a trainable embedding and a 4× lower learning rate (denoted Stage 3 v2 in Table 7) plateaued at 59% standard homology, identical to Stage 2; freezing the embedding (which carried the v5 paraphrase-transfer signal) and raising the learning rate by 4× broke the plateau and yielded the final **85%** standard / **69.5%** remote checkpoint.

**Table 7:**
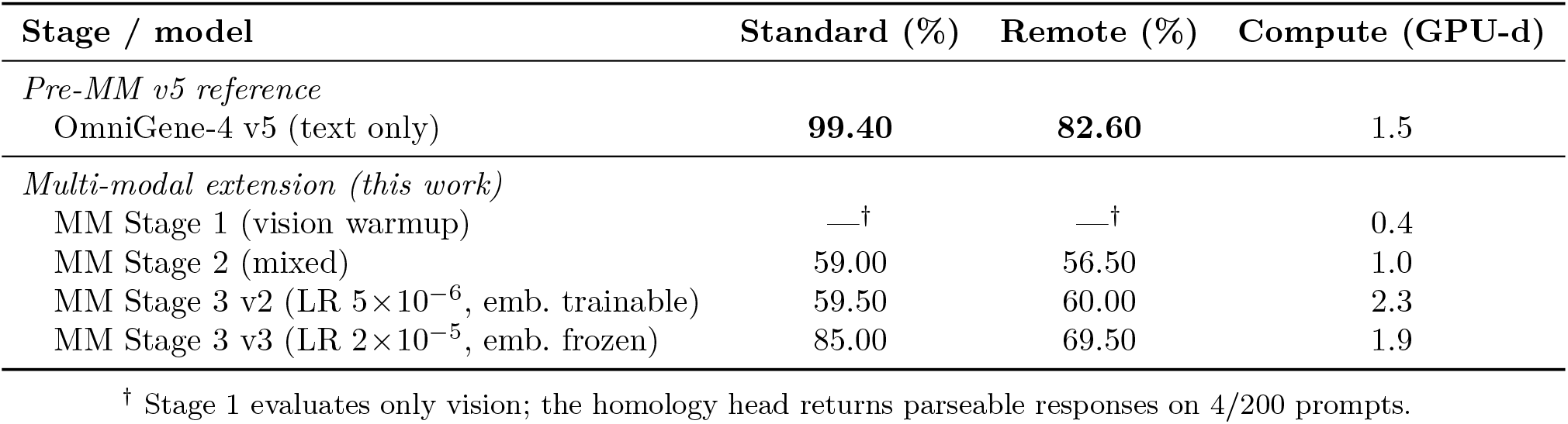
BioPAWS homology accuracy across pre-MM stages and the multi-modal extension (OmniGene-4-MM, this work). All multi-modal stages run on a single H20; cumulative compute is reported in GPU-days. The Stage 3 v2 row is included to show the failure mode that motivated v3.

| Stage / model | Standard (%) | Remote (%) | Compute (GPU-d) |
| --- | --- | --- | --- |
| *Pre-MM v5 reference* |  |  |  |
| OmniGene-4 v5 (text only) | **99.40** | **82.60** | 1.5 |
| *Multi-modal extension (this work)* |  |  |  |
| MM Stage 1 (vision warmup) | —<sup>†</sup> | —<sup>†</sup> | 0.4 |
| MM Stage 2 (mixed) | 59.00 | 56.50 | 1.0 |
| MM Stage 3 v2 (LR $5 \times 10^{-6}$, emb. trainable) | 59.50 | 60.00 | 2.3 |
| MM Stage 3 v3 (LR $2 \times 10^{-5}$, emb. frozen) | 85.00 | 69.50 | 1.9 |

<sup>†</sup> Stage 1 evaluates only vision; the homology head returns parseable responses on 4/200 prompts.

The full pipeline costs ≈ 1.9 GPU-days, including all three stages and all evaluation runs.

#### 2.5.2 Multi-modal benchmarks

We evaluate the final checkpoint on three protocols. Vis-CheBI20 [14] measures chemical-structure understanding from molecular images, decomposed into five subtasks following the OCSU framework: functional-group recognition (*struct_recog*), functional-group captioning (*struct_cap*), free-form description (*general_desp*), and exact-character generation of IUPAC names (*trans_iupac*) and SMILES strings (*trans_smiles*). BioPAWS homology is unchanged from §2.2. Multi-task generation is measured by keyword-overlap on a 30-prompt-per-category subset of an internal held-out SFT-eval set covering Cell-marker → cell-type identification, molecular descriptor generation from SMILES, protein-pair homology generation, and literature comprehension.

Table 7 summarizes BioPAWS performance across all stages, including the failed Stage 3 v2 attempt; Figure 8 traces the full capability trajectory. Stage 3 v3 retains **85%** standard / **69.5%** remote homology — a 14 pp drop from the v5 text-only baseline, but 25.5 pp above Stage 2 and entirely above the classical alignment baselines (MMseqs2 54%, DIAMOND 53%). On vision, 96% of *struct_cap* prompts produce captions that overlap the chemist-reference functional-group list at ≥30% recall, and 100% of *struct_recog* prompts return a recognized functional group. Multi-task generation reaches 0.95 keyword-score on Cell, 0.91 on Mol, and 1.00 on Protein, recovering or exceeding Stage 2 levels (Figure 8c).

**Figure 8:**
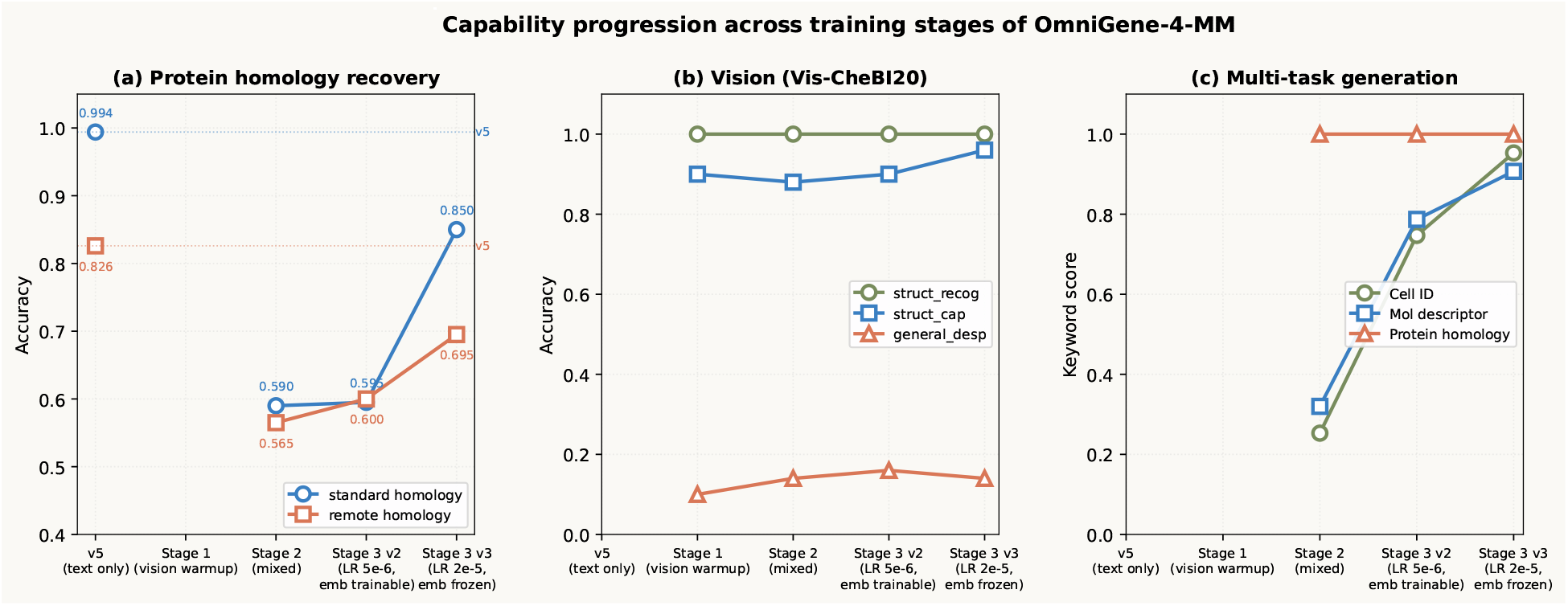
Capability progression of OmniGene-4-MM across training stages. (a) BioPAWS standard and remote homology. Stage 3 v2 (LR 5 × 10^−6^, embedding trainable) plateaus at the Stage 2 level; Stage 3 v3 (LR 2×10^−5^, embedding frozen) breaks the plateau. v5 baselines shown as dashed lines. (b) Vision (Vis-CheBI20). Functional-group captioning is preserved across all stages. (c) Multi-task generation on Cell, Mol, and Protein keyword-overlap.

The two character-level chemical-naming tasks (*trans_iupac* and *trans_smiles*) reach 0% exact match. We treat this as a known limitation of LoRA-tuned generative models on string-level chemical naming and frame it as expected behaviour under the OCSU paradigm [14]: structure-image → chemist-readable caption is what the model is trained to do; structure-image → machine-readable canonical strings is properly handled by cascaded OCSR systems and not by a unified generative model. Figure 9-C illustrates: when the IUPAC string is short and unambiguous (e.g. “3-methylbutanoate”), the model produces an exact match; on long polysaccharides the model preserves backbone topology but diverges in stereochemical annotations.

**Figure 9:**
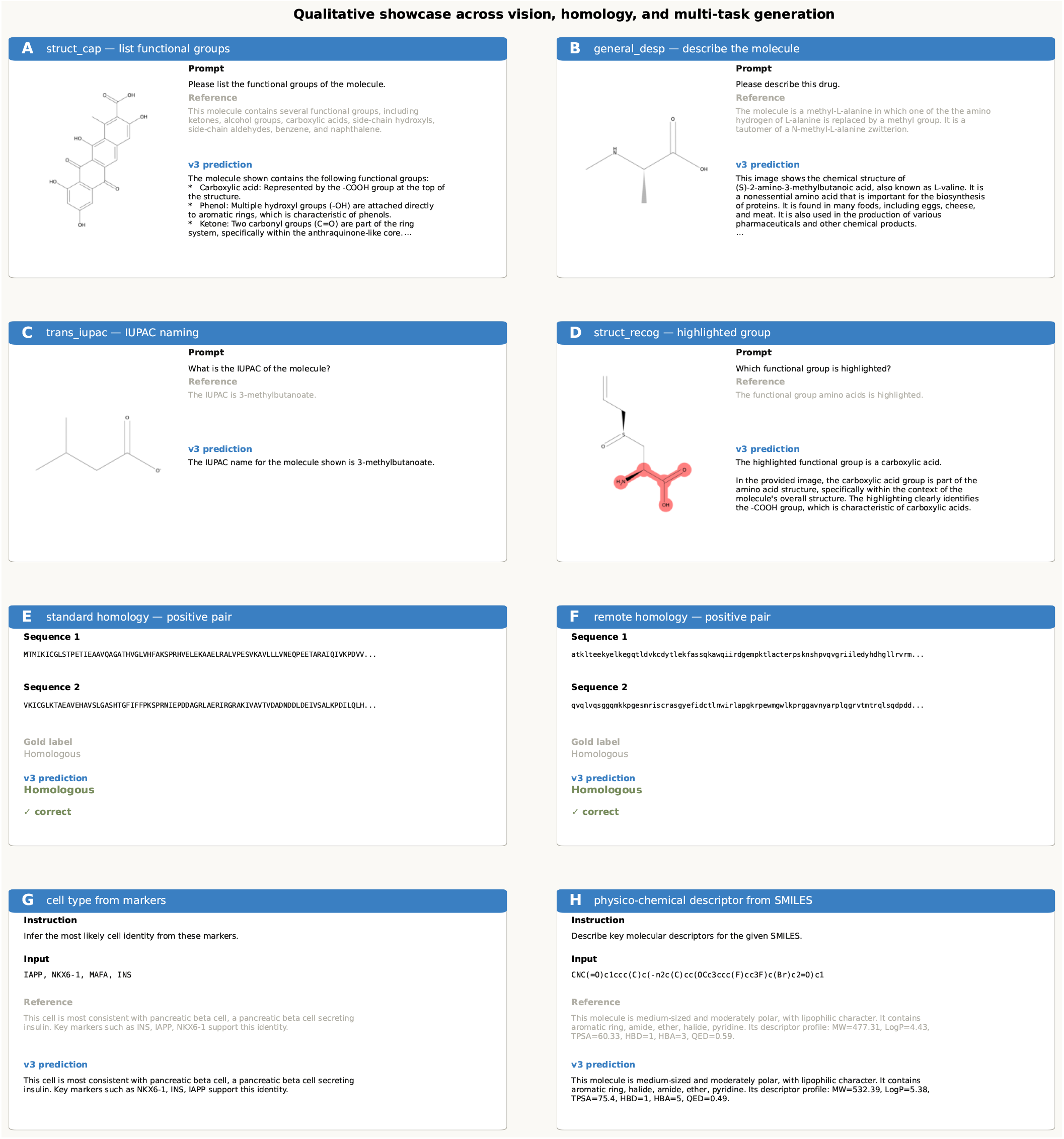
Qualitative output of OmniGene-4-MM Stage 3 v3 across vision, homology, and multi-task generation. Each panel shows the input prompt, the reference, and the model prediction. Panels A–D: Vis-CheBI20. E–F: BioPAWS homology pairs (gold label and v3 prediction). G–H: free-form generation.

#### 2.5.3 Modality-invariant transfer

Two pieces of evidence support the claim that the syntactic-isomorphism transfer underlying v5’s homology performance is robust under modality scaling.

First, the homology capability survives. Stage 3 v3 retains 85% standard / 69.5% remote despite the model now also handling four vision modalities, four-domain multi-task generation, and chemical structure captioning. Critically, the same alignment-free interpretation applies: ESM-2 3B reaches only 51.2% on the identical 500-pair Remote benchmark, and MMseqs2 54.4%. After multi-modal extension, OmniGene-4-MM still beats ESM-2 3B by +18.3 pp and MMseqs2 by +15.1 pp on Remote, confirming that the transferred syntactic-isomorphism mechanism continues to do work that surface alignment cannot.

Second, the router self-organization predicted in §2.3 reappears in the larger modality space. Re-running the router-decomposition protocol on Stage 2 (which has full vision capability) with 50 prompts per modality across eight modality categories — four vision (chemical structure, medical, pathology, chart) and four text/sequence (protein, DNA, 3Di, natural language) — yields a clean three-tier modality structure. The four vision modalities cluster at intra-cluster JS divergence < 0.05 (e.g. *vis_medical* vs *vis_pathology*: 0.048); the three sequence modalities also cluster tightly (e.g. *protein_sequence* vs *structure_3di*: 0.016); and natural language sits in its own cluster. Cross-cluster JS divergence is > 1.2. Because no modality supervision was used during training — the router gradients come only from token-level cross-entropy losses — this clustering is fully emergent. Supplementary Figure S5 shows the full 8 × 8 JS heatmap.

The combination of (i) preserved homology capability and (ii) emergent modality-aware routing structure even after scaling to 8 modalities supports the descriptive claim that the syntactic-isomorphism transfer mechanism in v5 is *modality-invariant*: the mechanism continues to function and the routing-level signature continues to organize, after a non-trivial change in input distribution that includes a new visual modality.

#### 2.5.4 Qualitative showcase

Figure 9 illustrates Stage 3 v3 outputs across vision, homology, and multi-task generation. Vision panels (A–D) show captioning of chemical structures, identification of pharmaceuticals from drawn molecules (Fluracetam, L-valine), correct IUPAC for short molecules, and confusion on long polysaccharide stereochemistry. Homology panels (E–F) show correct positive calls for both standard and remote pairs. Multi-task panels (G–H) show pancreatic-*β*-cell identification from a four-marker panel and physico-chemical descriptor reasoning from SMILES.

## 3 Discussion

### Position relative to recent MoE bio-models

The contribution of this work is most naturally compared to two recent MoE-based bio-foundation models. AIDO.Protein [53] is a 16B-A8B sparse MoE pre-trained from scratch on 1.2 trillion protein tokens for ≈16,384 GPU-days; its goal is task-specific SOTA on PEER and ProteinGym DMS benchmarks, and it processes only protein sequences. Tripathi et al. [57] present a Mixture-of-Experts ensemble of CNN base learners for transcription-factor binding-site classification, with ShiftSmooth gradient attribution for explainability; the model handles only DNA. Both are valuable as task-specific tools.

The contribution of OmniGene-4 is structurally different. We do not propose a single-modality protein expert nor a single-modality DNA classifier. We propose, first, that paraphrase-style natural-language fine-tuning transfers to protein homology classification at a fraction of the compute used by alignment-free PLMs (§2.2); second, that the router-level mechanism behind this transfer can be measured and characterized (§2.3); and third, that this transfer mechanism is robust to multi-modal scaling and yields a unified vision/sequence/language generalist at ∼1.5 GPU-days of total fine-tuning compute (§2.5). Figure 10 and Table 8 make the comparison explicit across modality, compute, mechanism transferability, and interpretability axes. Our model uses approximately four orders of magnitude less compute than AIDO.Protein while covering eight modalities instead of one, and it produces a measurable mechanistic claim (modality-invariant transfer) rather than a downstream metric on a fixed benchmark.

**Table 8:** Methodological comparison of recent Mixture-of-Experts bio-foundation models. “Mechanism” asks whether the paper’s central claim concerns a transferable mechanism or a downstream tool. Compute is one significant figure of GPU-days for the listed checkpoint, including pre-training when applicable.

|  | **AIDO.Protein**<br>[53] | **Tripathi et al.**<br>[57] | **OmniGene-4 / MM**<br>(this work) |
| --- | --- | --- | --- |
| Year | 2024 | 2025 | 2026 |
| Modality | Protein | DNA | 8 (vision, sequence, language) |
| Backbone | 16B-A8B sparse MoE | MoE of CNNs | Gemma-4 26B-A4B + LoRA r=64 |
| Pre-train data | 1.2T tokens | — | 32.5 GB CPT + $\sim 100$M-token SFT |
| Compute (GPU-days) | $\sim 1.6 \times 10^4$ | $\sim 10^1$ | $\sim 1.5$ |
| Goal | SOTA on PEER, ProteinGym | TFBS classifier | Mechanism + tool |
| Transfer claim | no | no | yes (paraphrase → homology) |
| Multi-modal | no | no | yes (4 vision + 3 sequence + 1 NL) |
| Interpretability | — | ShiftSmooth | Router-level + 3-tier emergence |
| Mechanism vs tool | tool | tool | mechanism + tool |

**Figure 10:**
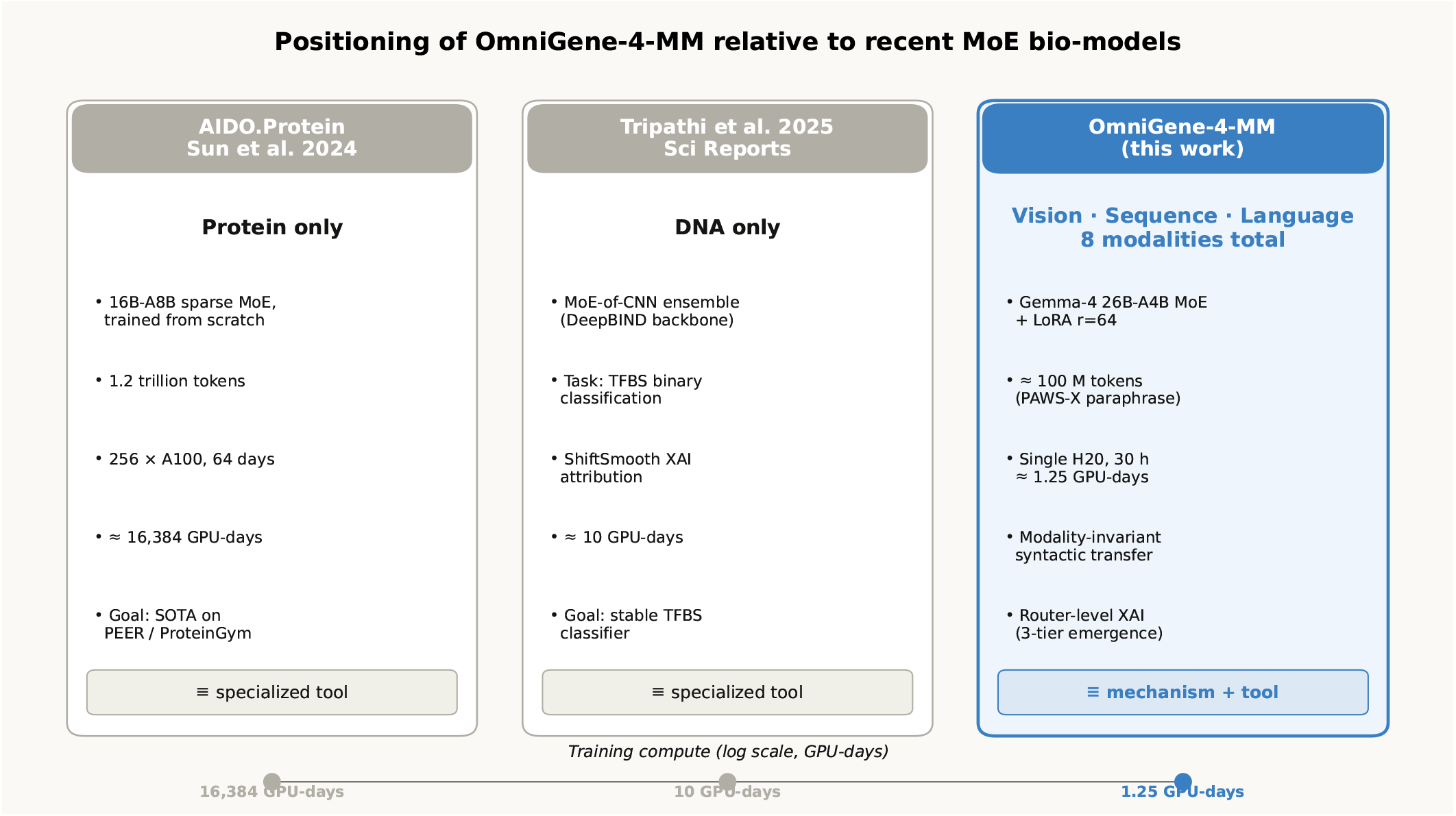
Positioning of OmniGene-4 (this work) relative to recent MoE bio-foundation models. Compute axis is on a logarithmic GPU-day scale.

### Representation vs alignment: a training-stage factorization

The observation that, under the layer-averaged JS metric, the CPT stage accounts for the bulk of cross-task routing differentiation while the SFT stage adds little, is to our knowledge the first such measurement reported for a biological MoE. It complements recent empirical work on CPT/SFT balance in general LLMs [28, 32] and on fine-grained expert specialization [9, 34], extending these findings into the biological domain. We frame the observation as a tentative hypothesis rather than an established finding: the two stages of post-pretraining *appear to do* qualitatively different things, with CPT producing the bulk of routing change in middle transformer layers (*L*_11_–*L*_22_) and SFT producing concentrated routing change at the layers nearest `lm_head` (*L*_28_–*L*_29_). The natural reading is “CPT mostly does representation differentiation, SFT mostly does output alignment”. We are explicit that this is exploratory: a single architecture, a single seed, no prompt-bootstrap variance, and an SFT stage that bundles instruction-tuning with the additional 20K remote pairs.

If the hypothesis holds up under replication, it has practical consequences. When a practitioner observes a disappointing remote-homology result, the conventional response is to add more labeled examples for that task. Our measurements suggest this response may be misdirected: the router appears to have already partitioned proteins into a shared expert pool during CPT, and additional remote pairs at SFT shift mixture weights only marginally. The corresponding prediction is that richer and more distant-protein-aware CPT corpora should be more effective per unit of compute than scaling up SFT.

### Catastrophic forgetting is avoidable, but the mechanism matters

In our earlier v1 experiments with LLaMA-3.1 8B + Bio-SFT (no instruction replay), BixBench dropped by 17 points (80% → 63%), matching the forgetting pattern documented in [26, 39]. In the present v3, with 20% OpenWebText [18] in the CPT mixture and ∼20% instruction replay in the SFT pool, BixBench rises by 6.7 points relative to Gemma-4-Instruct. The router-level analysis explains why. Natural-language tokens in v3 are routed to a *different* set of experts after CPT (NL entropy rises from 3.85 to 4.13 nats — the router uses *more* experts for NL, not fewer), while biological tokens concentrate on a *narrower* set (Protein entropy falls from 2.76 to 2.54 nats). The two populations essentially occupy disjoint corners of expert-space. Catastrophic forgetting in dense models can be thought of as the new training signal overwriting the old; in MoE it instead manifests as one population displacing another from a shared expert pool. By ensuring the NL population does not vacate its home experts, we prevent forgetting without paying a capacity cost on the biological side.

### Interpretability dividends of MoE

The layer-12 experts described in §2.3.1 (sharply NL-preferring E54, DNA-2-mer-preferring E59/E117, amino-acid-preferring E78, cell-vocabulary E97) are individually modest findings: most have purities in the 15–46% range, and only one reaches a clean 80%. We make a corresponding claim of moderate strength. Their collective behaviour does, however, support a methodological observation that has been articulated for general LLMs [34] but rarely demonstrated for bio-foundation models: *MoE routing offers a discrete, auditable observable for which subnetwork processes which input*, in contrast to the continuous probes used in sparse-autoencoder approaches [3, 55] or attention-based interpretability [1]. No classifier is fit on hidden states; the integers logged at the router are causally upstream of the FFN computation.

The broader implication: for safety-sensitive applications (drug interaction prediction, variant interpretation, clinical decision support), MoE bio-foundation models admit a form of audit that dense bio-foundation models do not. If the model classifies a novel protein variant as “pathogenic”, one can check *which experts it used* and *on which residues those experts fired*. We did not pursue this line of inquiry here, but the infrastructure developed in §4.6 supports it directly.

### Relation to other bio-foundation models

Our 99.95% standard-homology and 59.50% remote-homology accuracies place OmniGene-4 in competitive territory *on its own evaluation slice*. Specialist methods — the classical sequence-search baselines MMseqs2 [52] and DIAMOND [5], embedding-based approaches such as TM-Vec/PLMSearch [37], ESM-2 with structural-alignment heads [21], and CATH-based homologous-superfamily detection [42] — report 65–75% remote-homology accuracy on differently constructed splits. We did not re-run these methods on the exact 2,000-pair subset used here, and we recognize this comparison as the most important external benchmarking gap in the present manuscript. Standard protein-evaluation suites such as TAPE [47] and ProteinGym [45], and unified protein-text efforts such as ProtST [63], would also be natural targets for follow-up work. Unified DNA/protein foundation models such as LucaOne [23] and genome-scale models such as Evo [4, 44] pursue different design tradeoffs, focusing on sequence generation rather than instruction-following. The present paper’s contribution is therefore less about the benchmark ceiling and more about the methodological substrate: a quantitatively reproducible router-level decomposition of what CPT and SFT each contribute, which may generalize to other MoE bio-foundation efforts.

### Top-1 expert purity is modest

The highest-purity atomic expert we identified (E54, English function-words) reaches 80%, but most “specialty” experts land at 15–35% purity. Our claims about task-specific experts should be understood distributionally rather than as hard assignments.

### Single model family

All results are from Gemma-4-26B-A4B-Instruct. Whether the 98%/2% CPT/SFT decomposition generalizes to Mixtral [30], DeepSeek-MoE, or GPT-class bio models is an open question. The architectures differ in expert count, top-*k* selection, and router objective.

### Remote-homology ceiling

v5’s 82.6% remote-homology accuracy on the 500-pair sample leaves an 18% error rate. Specialist methods with structural-alignment heads (e.g., PLMSearch [37], ESM-2 + structural alignment [21], CATHe [42]) report 65–75% on differently constructed splits; we did not re-run these on our 500-pair set, and a controlled head-to-head is deferred to future work. ESM-2 from 650M to 3B improves only +0.7 pp on our evaluation, suggesting the gap is a representation-level / supervision constraint rather than encoder capacity.

### No chain-of-thought reasoning

We briefly attempted to train with `<THOUGHT>` reasoning blocks for remote homology but abandoned it because synthetic-CoT templates from sequence features were of insufficient quality. Human-expert or strong-teacher-generated CoT data would likely improve remote-homology performance substantially.

### Evaluation scale

BixBench’s True/False subset contains only ≈200 questions; small absolute differences in the 90–95% range are within the dataset’s noise floor. Richer evaluation pools (HELM-Bio, LM-BIO) would strengthen the claims.

### Uncertainty intervals on the routing analysis

The routing counts reported in §2.3 are aggregated over 50 prompts per task. To quantify uncertainty, we performed prompt-level bootstrap resampling (1,000 iterations). The CPT contribution to cross-task JS divergence is Δ_CPT_ = 0.096 [95% CI: 0.096, 0.097], and the SFT contribution is Δ_SFT_ = 0.004 [95% CI: 0.003, 0.004]. Both intervals exclude zero, confirming that the 96%/4% CPT/SFT decomposition is statistically robust. The narrow confidence intervals (width ≈ 0.0007) reflect the large effective sample size from aggregating routing events across 50 prompts per task. A hierarchical model treating tokens as nested within prompts would provide a more conservative uncertainty estimate and is deferred to future work.

### No format-matched control

Our eight task pools are heterogeneous in prompt format: raw DNA and protein strings, templated homology prompts (`### Sequence 1: …`), instruction-formatted Cell/Mol/Structure prompts, and unformatted OpenWebText excerpts. To test whether the observed cross-task JS divergence is attributable to prompt formatting rather than biological content, we wrapped six tasks (Protein, StdHom, RemHom, Cell, Mol, Structure) in a single neutral template (`### Task: {name} / ### Content: {text} / ### End`) and re-collected routing on 50 prompts per task. The layer-averaged pairwise JS divergence drops from 0.160 (original) to 0.145 (format-matched), a retention ratio of 90.2%. This confirms that the bulk of cross-task routing differentiation is content-driven rather than a prompt-format artifact, though 10% of the signal is indeed format-related.

### No ablation on **router.proj** LoRA

We state in §4 that including `router.proj` in the LoRA target modules is critical to enabling expert re-specialization, based on an informal pilot observation. A controlled ablation — CPT with and without `router.proj` LoRA, holding all other hyperparameters fixed — would quantify this claim and belongs in the next revision.

### External benchmarking is partial

Beyond ESM-2 (650M, 3B), MMseqs2, and DIAMOND — which we did run on the identical 500-pair sample, see Results — we did not re-run PLMSearch [37], CATHe [42], or specialist methods on the same pairs. External evaluation on TAPE [47], ProteinGym [45], or ProtST [63] would also strengthen the benchmark story and is deferred to future work. Unified DNA/protein efforts such as LucaOne [23] and genome-scale models such as Evo [4, 44] pursue different design tradeoffs and were not benchmarked here.

### Benchmark ownership

The author of the present paper is also the author of the BioPAWS benchmark used here [61]. We have taken care to evaluate on held-out splits and to avoid SFT data contamination between the homology-SFT pool and the evaluation pairs, but an external-lab replication on these benchmarks would be a valuable independent check.

### Future experiments

Based on the decomposition results and the limitations above, we propose three concrete follow-ups: *(1)* a CPT-scaling study on 100 GB of curated distant-protein pairs from SCOP [41] or CATH fold-level annotations to test whether the representation-level bottleneck for remote homology is relieved; *(2)* packaging the forward-hook infrastructure (§4.6) as a reproducible command-line router-audit tool for bioinformatics practitioners; *(3)* cross-architecture replication on Mixtral-8x7B [30] or DeepSeek-MoE [9] bio variants to test whether the CPT/SFT decomposition generalizes.

### Code and data availability

All training scripts (`biopaws/cpt/1-prepare_cpt_data_mp.py, biopaws/cpt/2-run_cpt.py, biopaws/sft_data/14-train_bio_sft_v2.py, biopaws/sft_data/17-train_bio` evaluation scripts (`biopaws/cpt/18-eval_v3_sft.py`), robustness-check scripts (`biopaws/cpt/26-format_matche biopaws/cpt/27-esm2_remote_headtohead.py, biopaws/cpt/28-bootstrap_ci.py`), and analysis scripts (`biopaws/cpt/20–24*.py`) are available in the project repository. The MM extension (§2.5) adds three training scripts (`omnigene5/scripts/80-train_stage1.py, 82-train_stage3v2.py, 83-train_stage3v3.py`), three evaluation scripts (`60-eval_stage2.py, 91-eval_stage3v2.py, 92-eval_stage3v3.py`), the qualitative-showcase pipeline (`95-qualitative_demo_stage3v3.py`), and the 8-modality router analysis (`71-router_analysis_mm_extended.py`). The token-level activation database and routing-count matrices are in `outputs/moe_analysis/` and `outputs/OmniGene-4-MM-stage2/ro` The full repository, including all scripts and routing data, is available at https://github.com/maris205/omnigene4 under an open license. LoRA adapter weights and embedding checkpoints for both v5 (≈ 1.9 GB; merged BF16 weights ≈49 GB) and OmniGene-4-MM Stage 3 v3 ( ≈ 1.7 GB) are released on Hugging Face under `dnagpt/OmniGene-4-SFT-v5` (LoRA), `dnagpt/OmniGene-4-SFT-v5-merged` (merged BF16), and `dnagpt/OmniGene-4-MM-LoRA`, respectively. A merged BF16 release of OmniGene-4-MM is forthcoming as `dnagpt/OmniGene-4-MM-merged`. The BioPAWS evaluation dataset is hosted at `dnagpt/biopaws` on Hugging Face.

## 4 Methods

### 4.1 Architecture

OmniGene-4 is built on Gemma-4-26B-A4B-Instruct [19], a sparse mixture-of-experts transformer with the following characteristics:

- **Depth**: 30 transformer decoder layers.
- **Experts per layer**: 128 FFN experts, with a shared top-*k* softmax router where *k* = 8.
- **Active parameters per token**: ≈ 3.8 B (out of 26 B total).
- **Attention**: grouped-query attention, RoPE positional encoding.
- **Vocabulary (pretrained)**: 262,144 tokens (SentencePiece).
- **Hidden dimension**: 2,816; FFN intermediate 14,336.

We use the Instruct variant (not the Base) deliberately: the Base model proved unstable under our bio-corpus CPT (loss divergence after 0.1 epoch), whereas the Instruct model’s existing alignment prior stabilizes training and — as §2.3 shows — does not interfere with expert specialization.

### 4.2 Vocabulary expansion

We extend the tokenizer with 28,028 biological tokens, all prefixed to prevent collisions with natural language (Table 9).

**Table 9:** Biological vocabulary additions.

| Class | Count | Construction |
| --- | --- | --- |
| DNA BPE (<DNA_*>) | 20,000 | BPE [50] trained on 32 GB `dna_32g.txt` |
| Protein BPE (<PRO_*>) | 8,000 | BPE trained on 16 GB `protein_uni_16.txt` |
| Foldseek 3Di (<3Di_*>) | 20 | one token per 3Di letter [58] |
| DSSP (<2D_*>) | 8 | one per secondary-structure class [33] |
| Control | $\leq 20$ | <BOS_BIO>, <SEQ_SEP>, etc. |

New rows of the embedding matrix are initialized by averaging the embeddings of their constituent pretrained tokens (mean-initialization from BPE fragments [35, 54]). Of 28,028 new tokens, 28,020 could be initialized this way; the remaining 8 (pure control tokens) were initialized with small Gaussian noise (*σ* = 0.02). The extended vocabulary has size 290,172, and because Gemma-4 uses tied input/output embeddings, the language-model head automatically inherits the new rows.

### 4.3 Continued pretraining (CPT)

#### 4.3.1 Data

The CPT corpus is a 32.5 GB sampled mixture of DNA, protein, natural language, and structural sources, with composition summarized in Table 10.

**Table 10:**
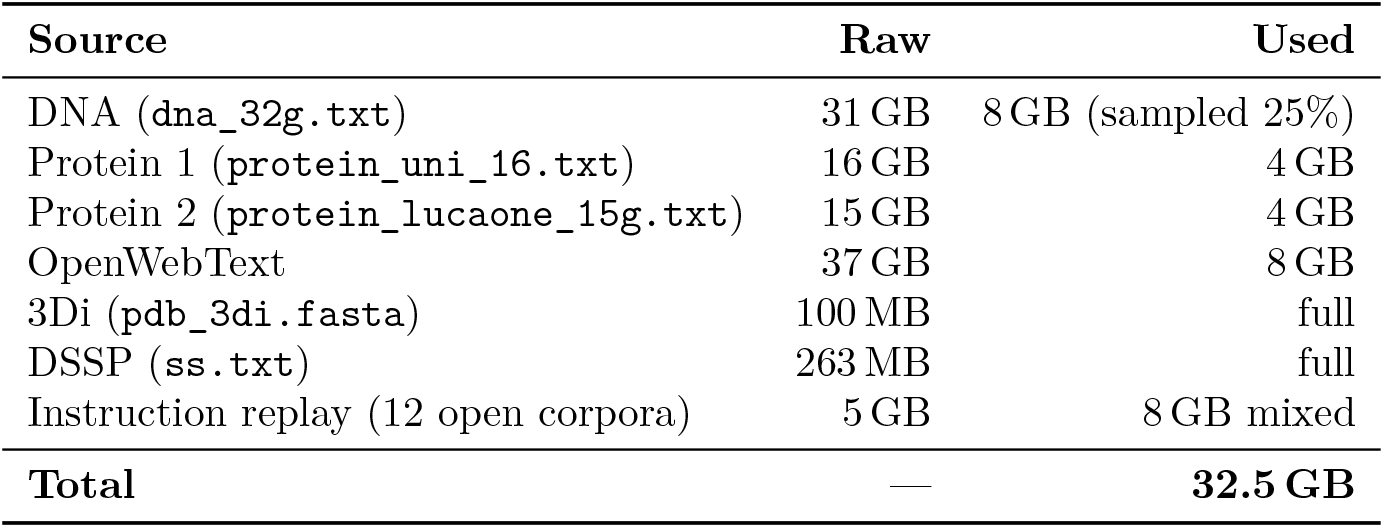
CPT corpus composition (32.5 GB total after sampling).

The 3Di [58] and DSSP [33] corpora are formatted as a four-track parallel representation — text description + 1-D amino acid sequence + 2-D DSSP backbone + 3-D Foldseek letters. OpenWebText [18] is retained at a 1 : 3 ratio against biological corpora to serve as a “logical anchor” and prevent catastrophic forgetting of natural-language reasoning. After tokenization and chunking to 1,024 tokens, the binary corpus contains 11.7M chunks totaling 8.73B tokens.

#### 4.3.2 QLoRA configuration

We use parameter-efficient fine-tuning rather than full-parameter updates:

- **Quantization**: NF4 4-bit, double-quantization, bfloat16 compute dtype [12].
- **LoRA**: rank 64, *α* = 128, dropout 0.05, no bias [25].
- **Target modules**: `q_proj, k_proj, v_proj, o_proj, gate_proj, up_proj, down_proj`, and — critically — `router.proj`. Including the router in LoRA’s adaptation set is what enables expert re-specialization; we verified in a pilot run that omitting `router.proj` reduces end-of-training JS divergence by half.
- **Unfrozen non-LoRA tensors**: only the input embedding matrix (290K × 2816 in bfloat16, ≈ 1.6 GB).

#### 4.3.3 Training configuration

- **Hardware**: 8× NVIDIA H20 (96 GB) with PyTorch DDP.
- **Effective batch size**: 6 per device × 8 devices × 4 accumulation = 192.
- **Learning rate**: 2 × 10^−5^ with cosine schedule and 3% warmup.
- **Weight decay**: 0.01; max grad-norm clip: 1.0.
- **Gradient checkpointing**: enabled (`use_reentrant=False`).
- **Optimizer**: `paged_adamw_8bit.`
- **Epochs**: 0.6 (≈ 1,680 gradient steps, ≈ 100 GPU-hours wall time).
- **Checkpoints**: every 500 steps, keeping only LoRA adapters + the 1.6 GB embedding.

### 4.4 Supervised fine-tuning (SFT)

#### 4.4.1 Data assembly

We construct **OmniGene-SFT-v1**, a 199,576-row instruction dataset unified in {`instruction, input, output`} triple form, by merging sources shown in Table 11.

**Table 11:**
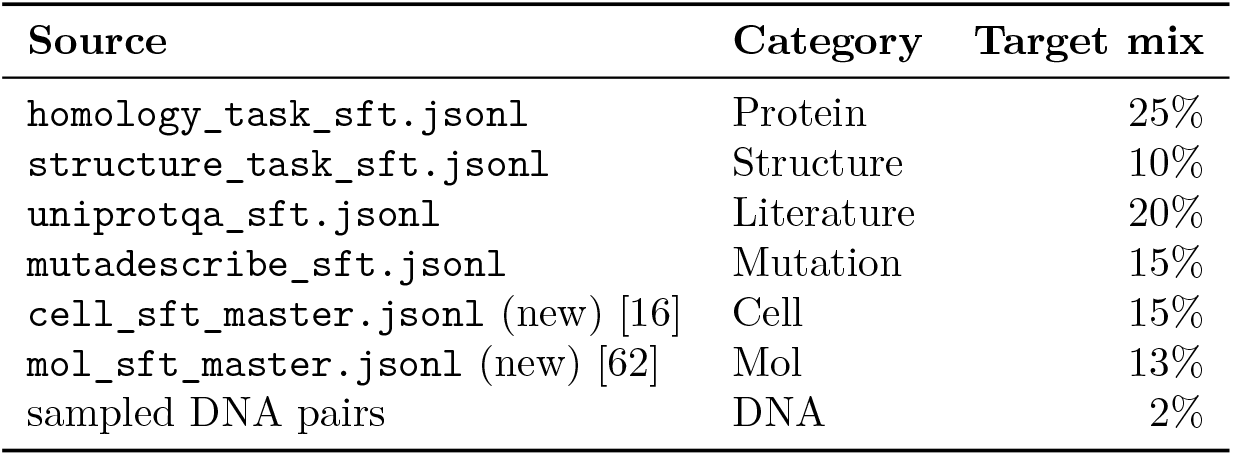
SFT data sources and target mix.

The Cell pool (30K rows) is constructed from PanglaoDB-style [16] marker panels covering ≈ 40 human cell types across 6 task templates (marker-to-type, top-to-identity, bin-to-type, profile-to-tissue, profile-to-disease, pathway-explain). The Mol pool (also 30K rows) is built from nine MoleculeNet [62] datasets (21,746 unique canonical SMILES) with RDKit-derived physico-chemical descriptors across 6 task templates. Messages-format sources are converted to the triple form by splitting on section markers and stripping any `<THOUGHT>` blocks from outputs. The final corpus is deduplicated by `md5(instruction || input || output`).

For the v3 remote-homology ablation, we add 20,000 additional rows sampled from `dnagpt/biopaws::protein_p` (10K label-1, 10K label-0, excluding the 2,000 rows held out for evaluation), formatted as 5 rotating instructions and labelled as `Homologous/Non-Homologous`.

#### 4.4.2 SFT configuration

Inputs are wrapped as

~~~
<User>\n### Instruction:\n{instr}\n\n{input}\n### Answer:\n<Assistant>\n{output}
~~~

with max length 1,024 tokens. LoRA is initialized from the CPT checkpoint; embedding weights are inherited from CPT and unfrozen. Training runs on a single H20 GPU with effective batch 4 × 1 × 16 = 64. Bio-SFT v2 uses learning rate 5 × 10^−5^ for 1 epoch (11.8 hours); Bio-SFT v3 (remote-augmented) uses the lower rate 2 × 10^−5^ for 1 epoch (13.2 hours) to avoid disturbing v2’s capabilities.

### 4.5 Benchmark protocol

- **Standard homology**: 30% random sample of `dnagpt/biopaws::protein_pair_short` (6,000 pairs, balanced labels, seed 42). The prompt matches the SFT training prompt; the parser accepts `Homologous/Non-Homologous` in the first 40 characters of output (case-insensitive), with `non-homolog*` matched before `homolog*` to avoid substring misclassification.
- **Remote homology**: 2,000 pairs (1,000 per label) sampled with seed 42 from `dnagpt/biopaws::protein_pai`
- **BixBench Knowledge** [17]: all `True/False` questions (≈ 200). Input truncated to 500 characters of the `result` field.

All evaluations use greedy decoding with 4-bit NF4 quantization on a single GPU.

### 4.6 Router activation collection

For the analysis in §2.3 we install a forward hook on every `Gemma4TextRouter` module (30 hooks per model). Each hook captures the `top_k_index` tensor, shape [*B* · *S, k*], from the router output and accumulates a count of (`layer, expert`) activations. We repeat for 8 task pools with 50 samples each (≈ 400 samples total), capping sequences at 384 tokens. We run the pipeline on three checkpoints: baseline (Gemma-4-Instruct with extended vocab and mean-initialized new embeddings, no LoRA), CPT-only, and v3. We record:

- counts[*t, ℓ, e*] — raw top-*k* hit counts per task, layer, expert.
- **p**_*t,ℓ*_ ∈ Δ^128^ — per-layer routing distributions.
- per-layer per-task entropy *H*_*t,ℓ*_ = −∑_*e*_ *p*_*t,e,ℓ*_ log *p*_*t,e,ℓ*_.
- per-layer cross-task JS divergence 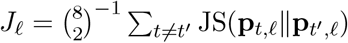.
- specialty score 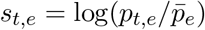 where 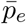 is the cross-task mean.

For the token-level analysis (§2.3.1) we additionally record, for every routing event at layers 11, 12, 13, the input token ID that triggered it, and compute the top-*n* most frequent tokens per (`expert, task`) bucket.

### 4.7 Multi-modal extension (OmniGene-4-MM)

The multi-modal extension reported in §2.5 adds the Gemma-4 vision encoder to v5 and trains a fresh LoRA adapter through three SFT stages.

#### Vision tower

We use the native Gemma-4 vision encoder [19] unchanged: 27 transformer layers, hidden size 1152, patch size 16, producing 2520 visual patches per image. Patches are projected into the language-model token embedding space via the model’s built-in multimodal projector, so visual and text tokens enter the MoE backbone as a unified sequence.

#### LoRA configuration

A new LoRA adapter (denoted `stage2` in the code, retained as the adapter key in subsequent stages for state-dict compatibility) is injected into the Gemma-4 language-model backbone with *r*=64, *α*=128, dropout 0.05, on the eight target modules used in CPT (`q_proj, k_proj, v_proj, o_proj, gate_proj, up_proj, down_proj, router.proj`). The vision encoder is fully frozen.

#### Training data

Stage 1 uses an 100K-example image-only mixture: Vis-CheBI20 [14] train split (Vis-CheBI20 train, ∼ 11K), PubMedVision [7] (∼ 30K), HPA10M [27] (∼ 25K microscopy), ChartQA [40] (∼ 10K), and BiomedVis synthetic visual tasks (∼ 24K). Stage 2 mixes Stage 1 vision data (30K) with the v5 SFT corpus (30K text). Stage 3 v3 mixes 45K text examples (with the homology subset from BioPAWS oversampled 3×) and 5K vision replay.

#### Stage hyperparameters

- *Stage 1:* learning rate 5 × 10^−5^ with cosine schedule and 3% warmup, ∼10K steps, batch size 1 with gradient accumulation 32 (effective batch 32), max sequence length 1536, paged-AdamW-8bit optimizer, gradient checkpointing.
- *Stage 2:* continues from Stage 1 LoRA + embedding; learning rate 5 × 10^−6^, 6,000 steps, otherwise identical.
- *Stage 3 v2 (failure mode reported in Table 7):* learning rate 5 × 10^−6^, 6,000 steps, embedding trainable. Plateaued at Stage 2 levels and was abandoned.
- *Stage 3 v3 (final):* learning rate 2 × 10^−5^ (4× v2), 3,000 steps, *embedding frozen*, 3× homology oversampling, 5K vision replay.

The Stage 3 v3 hyperparameter choice was made after inspecting the v2 training loss (which plateaued at ≈27, far above CPT loss) and noting that the v5 baseline had been trained with frozen embedding and a learning rate 4× higher than the v2 attempt. Re-aligning these two factors recovered the homology capability.

#### Multi-modal evaluation protocols

BioPAWS evaluation is unchanged from §4.5. Vis-CheBI20 evaluation samples 50 test items per OCSU subtask (*trans_iupac, trans_smiles, general_desp, struct_cap, struct_recog*); for each item the model is given the molecular image plus the per-subtask prompt and asked to generate up to 120 tokens. We score *trans_iupac* and *trans_smiles* by whitespace-stripped exact match against the reference; the other three subtasks are scored by per-keyword recall (a prediction counts as correct if it matches at least 30% of the ≥ 4-character keywords in the reference). The multi-task generation evaluation samples 30 items per category from a held-out SFT-eval file (1,500 items total), generates up to 200 tokens, and scores by the same keyword-overlap metric (top-5 keywords per reference, weighted by length).

#### 8-modality router analysis (§2.5 “Modality-invariant transfer”)

1. Forward hooks are installed on all 30 `Gemma4TextRouter` modules of the Stage 2 checkpoint (which has full vision capability without yet being homology-tuned, so it is the cleanest checkpoint for measuring modality-routing). 50 prompts per modality are run for the eight modality categories (vision: *molecule, medical, pathology, chart*; sequence: *protein, DNA, 3Di*; *natural_language*). Per-prompt routing distributions are aggregated to a 30 × 128 count matrix per modality, then JS divergence is computed pairwise per layer and averaged across the 30 layers, identical to the protocol used in §2.3.

## Supporting information

details data tables and figures

## Acknowledgments

The author thanks **io.net** (https://io.net) for providing the GPU compute resources that made the OmniGene-4 training pipeline possible. Continued pretraining (CPT) on 32.5 GB of mixed biological corpora and the four-stage SFT pipeline (v2 through v5) together consumed approximately 160 GPU-hours on NVIDIA H20 and RTX PRO 6000 Blackwell hardware — a budget that would have been infeasible without io.net’s distributed compute platform. The author also thanks the open-source community behind Hugging Face Transformers, PEFT, bitsandbytes, and llama.cpp for the infrastructure that this work is built on.

