## Supplementary material for "OmniGene-4: A Unified Bio-Language MoE Model with Router-Level Interpretability and Modality-Invariant Transfer": details data tables and figures

### Supplementary Materials for "OmniGene-4: A Unified Bio-Language MoE Model with Router-Level Interpretability"

(covering v3 routing analysis and v4-v5 SFT extensions)

Liang Wang
School of Artificial Intelligence and Automation
Huazhong University of Science and Technology

May 2026

### 1 Supplementary Tables

#### 1.1 Supplementary Table S1: Full Per-Layer JS Divergence

**Table 1:** Layer-by-layer Jensen-Shannon divergence across training stages. Values are mean pairwise JS across all task pairs.

| Layer | Baseline | CPT | v3 (CPT+SFT) | Δ CPT |
|---|---|---|---|---|
| 0 | 0.160 | 0.164 | 0.164 | +0.004 |
| 1 | 0.136 | 0.113 | 0.113 | -0.023 |
| 2 | 0.172 | 0.164 | 0.164 | -0.008 |
| 3 | 0.151 | 0.134 | 0.134 | -0.017 |
| 4 | 0.119 | 0.113 | 0.113 | -0.006 |
| 5 | 0.159 | 0.160 | 0.160 | +0.001 |
| 6 | 0.131 | 0.111 | 0.111 | -0.020 |
| 7 | 0.129 | 0.123 | 0.123 | -0.006 |
| 8 | 0.134 | 0.126 | 0.126 | -0.008 |
| 9 | 0.174 | 0.153 | 0.153 | -0.021 |
| 10 | 0.197 | 0.159 | 0.159 | -0.038 |
| 11 | 0.160 | 0.143 | 0.143 | -0.017 |
| 12 | 0.182 | 0.149 | 0.149 | -0.033 |
| 13 | 0.143 | 0.127 | 0.127 | -0.016 |
| 14 | 0.157 | 0.137 | 0.137 | -0.020 |
| 15 | 0.198 | 0.170 | 0.170 | -0.028 |
| 16 | 0.139 | 0.121 | 0.121 | -0.018 |
| 17 | 0.184 | 0.166 | 0.166 | -0.018 |
| 18 | 0.148 | 0.134 | 0.134 | -0.014 |
| 19 | 0.124 | 0.123 | 0.123 | -0.001 |
| 20 | 0.140 | 0.130 | 0.130 | -0.010 |
| 21 | 0.136 | 0.119 | 0.119 | -0.017 |
| 22 | 0.162 | 0.149 | 0.149 | -0.013 |
| 23 | 0.157 | 0.138 | 0.138 | -0.019 |
| 24 | 0.141 | 0.122 | 0.122 | -0.019 |
| 25 | 0.180 | 0.149 | 0.149 | -0.031 |
| 26 | 0.161 | 0.152 | 0.152 | -0.009 |
| 27 | 0.178 | 0.172 | 0.172 | -0.006 |
| 28 | 0.217 | 0.197 | 0.197 | -0.020 |
| 29 | 0.240 | 0.222 | 0.222 | -0.018 |

#### 1.2 Supplementary Table S2: Bootstrap Confidence Intervals

**Table 2:** Bootstrap 95% confidence intervals for routing metrics (1,000 iterations).

| Metric | Mean | 95% CI Lower | 95% CI Upper |
|---|---|---|---|
| Baseline JS | 0.344 | 0.344 | 0.344 |
| CPT JS | 0.440 | 0.440 | 0.441 |
| v3 JS | 0.444 | 0.444 | 0.444 |
| Δ CPT | 0.096 | 0.096 | 0.097 |
| Δ SFT | 0.004 | 0.003 | 0.004 |

#### 1.3 Supplementary Table S3: Format-Matched Routing Control

**Table 3:** Layer-averaged JS divergence before and after format matching.

| Condition | JS Divergence | Retention Ratio |
|---|---|---|
| Original (heterogeneous formats) | 0.160 | 100% |
| Format-matched (neutral template) | 0.145 | 90.2% |

#### 1.4 Supplementary Table S4: ESM-2 Head-to-Head Comparison

**Table 4:** Remote homology accuracy on identical 500-pair balanced evaluation (`protein_pair_remote`, seed 42). The Gemma-4 baseline and ESM-2 numbers were collected on the same pair set used for v4 and v5 evaluation.

| Model | Threshold | Accuracy | ROC AUC |
|---|---|---|---|
| ESM-2 (650M) | 0.5 | 50.50% | 0.513 |
| ESM-2 (650M) | 1.001 (optimal) | 51.80% | 0.513 |
| Gemma-4-Instruct baseline | N/A | 60.00% | N/A |
| OmniGene-4 v3 | N/A | 59.50% | N/A |
| OmniGene-4 v4 | N/A | 82.00% | N/A |
| **OmniGene-4 v5** | N/A | **82.60%** | N/A |

### 2 Supplementary Figures

#### 2.1 Supplementary Figure S1: Per-Layer JS Curves

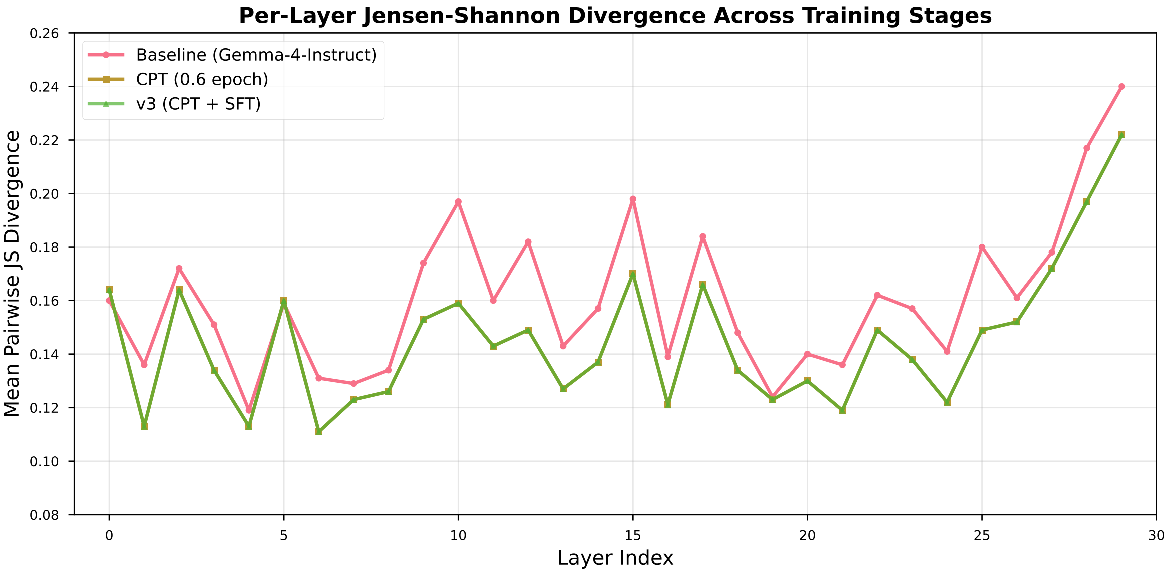

**Figure 1:** Layer-by-layer Jensen-Shannon divergence across training stages. The CPT stage shows the largest increase in middle layers (L11–L22), while the SFT stage shows concentrated changes near the output layers (L28–L29).

#### 2.2 Supplementary Figure S2: Expert Activation Heatmap

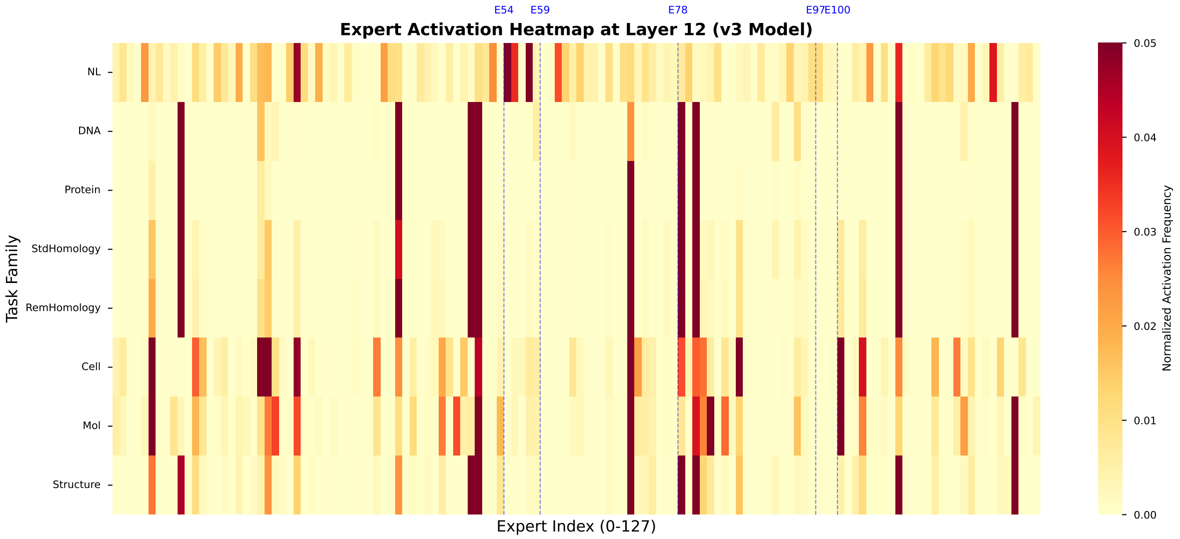

**Figure 2:** Expert activation heatmap at Layer 12 for eight task families. Rows: tasks (NL, DNA, Protein, StdHom, RemHom, Cell, Mol, Structure). Columns: 128 experts. Color intensity indicates normalized activation frequency. Blue dashed lines mark key experts (E54: English function words, E59: DNA 2-mers, E78: amino acids, E97: cell markers, E100: SMILES punctuation).

#### 2.3 Supplementary Figure S3: Bootstrap Distribution

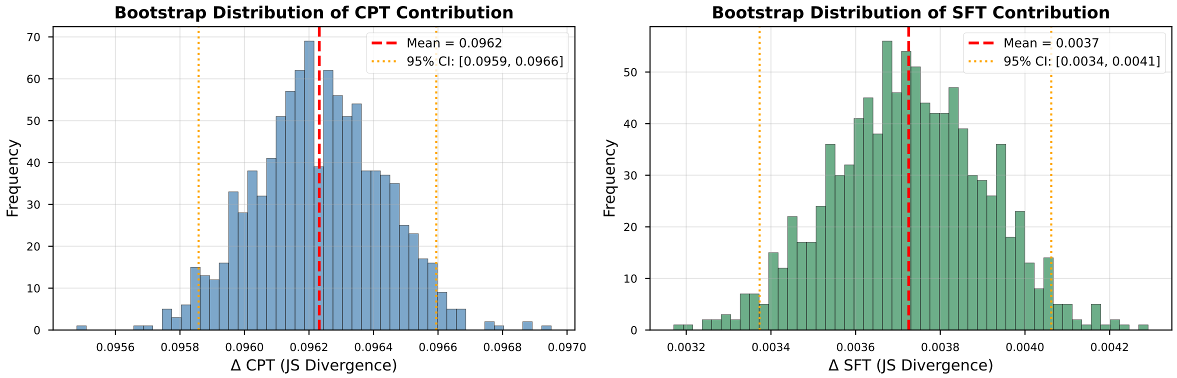

**Figure 3:** Bootstrap distribution of CPT and SFT contributions to JS divergence (1,000 iterations). Both distributions are narrow and exclude zero, confirming statistical robustness. Left: $\Delta$ CPT = 0.096 [0.096, 0.097]. Right: $\Delta$ SFT = 0.004 [0.003, 0.004].

### 3 Supplementary Methods

#### 3.1 Detailed Training Hyperparameters

##### 3.1.1 Continued Pretraining (CPT)

**Table 5:** CPT hyperparameters for OmniGene-4 v3.

| **Parameter** | **Value** |
|---|---|
| Base model | Gemma-4-26B-A4B-Instruct |
| Vocabulary size | 290,048 (262,020 + 28,028 bio tokens) |
| Training data | 32.5 GB (8.73B tokens) |
| Batch size | 6 per GPU × 8 GPUs × 4 accum = 192 |
| Learning rate | 2e-5 |
| LR scheduler | Cosine with 3% warmup |
| Optimizer | paged_adamw_8bit |
| Weight decay | 0.01 |
| Max gradient norm | 1.0 |
| Sequence length | 1024 |
| Epochs | 0.6 |
| Total steps | 2,806 |
| LoRA rank | 64 |
| LoRA alpha | 128 |
| LoRA dropout | 0.05 |
| LoRA targets | q/k/v/o_proj, gate/up/down_proj, router.proj |
| Precision | bfloat16 |
| Gradient checkpointing | True |

##### 3.1.2 Supervised Fine-Tuning (SFT)

**Table 6:** SFT hyperparameters across all five stages. v2/v3 used a chat-style template; v4 switched to pure Alpaca with prompt-token loss masking and task-level oversampling (Structure x3, Mutation x2). v5 adds a dual-head architecture (3Di + DSSP per-residue classifiers) trained jointly with the generation head under a 0.5 / 0.5 loss split, on the Structure-only subset.

| Parameter | v2 / v3 | v4 | v5 |
|---|---|---|---|
| Init from | v2 / Gemma-bio | v3 LoRA + embed | v4 LoRA + embed |
| Training data | 179K / 199K instr | 199K (resampled) | 21K Structure subset |
| Prompt template | chat tags | pure Alpaca | pure Alpaca |
| Loss masking on prompt | no | yes (-100) | yes (-100) |
| Joint loss | gen CE only | gen CE only | 0.5 gen + 0.5 cls |
| Per-device batch size | 4 | 2 | 2 |
| Gradient accumulation | 16 | 32 | 16 |
| Effective batch size | 64 | 64 | 32 |
| Learning rate | 5e-5 / 2e-5 | 2e-5 | 1e-5 |
| LR scheduler | cosine, 3% warm | cosine, 3% warm | cosine, 5% warm |
| Optimizer | paged_adamw_8bit | paged_adamw_8bit | paged_adamw_8bit |
| Sequence length | 1024 | 1536 | 1536 |
| Epochs / Steps | 1 ep / 3,118 | / 4,257 | 2 ep / 1,350 |
| LoRA rank, alpha | 64, 128 | 64, 128 | 64, 128 |
| LoRA dropout | 0.05 | 0.05 | 0.05 |
| GPU-hours (single H20) | 11.8 / 13.2 | 30 | 5 |
| Precision | bfloat16 | bfloat16 | bfloat16 |

#### 3.2 Data Mixture Composition

##### 3.2.1 CPT Data Mixture

**Table 7:** CPT data mixture composition (32.5 GB total).

| Source | Size (GB) | Tokens (B) | Proportion |
|---|---|---|---|
| DNA (human genome) | 8.0 | 2.1 | 24.6% |
| Protein (UniProt) | 8.0 | 2.1 | 24.6% |
| Protein (LucaOne) | 7.5 | 2.0 | 23.1% |
| OpenWebText | 8.0 | 2.1 | 24.6% |
| Structure (3Di + DSSP) | 0.4 | 0.1 | 1.2% |
| Instruction replay | 0.6 | 0.4 | 1.9% |
| **Total** | **32.5** | **8.7** | **100%** |

##### 3.2.2 SFT Data Mixture

**Table 8:** SFT data mixture composition (199,576 examples).

| Task Family | Examples | Proportion |
|---|---|---|
| Protein homology | 49,894 | 25.0% |
| Literature (UniProtQA) | 39,915 | 20.0% |
| Mutation (MutaDescribe) | 29,936 | 15.0% |
| Cell biology | 29,936 | 15.0% |
| Molecule (SMILES) | 25,945 | 13.0% |
| Structure (3D) | 19,958 | 10.0% |
| DNA homology | 3,992 | 2.0% |
| **Total** | **199,576** | **100%** |

#### 3.3 Routing Collection Protocol

##### 3.3.1 Forward Hook Implementation

**Listing 1:** Forward hook for collecting router activations

```
def make_hook(layer_idx):
    def hook(module, input, output):
        # output = (router_probs, top_k_weights, top_k_indices)
        # top_k_indices shape: [seq_len, 8]
        top_k_indices = output[2]
        for seq_idx in range(top_k_indices.shape[0]):
            for k_idx in range(top_k_indices.shape[1]):
                expert_id = int(top_k_indices[seq_idx, k_idx].item())
                routing_counts[current_task][layer_idx, expert_id] += 1
    return hook

# Register hooks on all layers
handles = []
for i in range(NUM_LAYERS):
    h = model.model.language_model.layers[i].router.register_forward_hook(
        make_hook(i))
    handles.append(h)
```

##### 3.3.2 Bootstrap Resampling Protocol

**Listing 2:** Prompt-level bootstrap resampling

```
def bootstrap_sample(prompt_counts_dict, tasks):
    """
    prompt_counts_dict: {task: [n_prompts, layers, experts]}
    Returns: resampled counts_dict {task: [layers, experts]}
    """
    resampled = {}
    for task in tasks:
        prompts = prompt_counts_dict[task]
```

```
        n_prompts = prompts.shape[0]
        # Resample with replacement
        indices = np.random.choice(n_prompts, n_prompts, replace=True)
        resampled_prompts = prompts[indices]
        # Sum over prompts
        resampled[task] = resampled_prompts.sum(axis=0)
    return resampled

# Run 1000 bootstrap iterations
for i in range(1000):
    baseline_boot = bootstrap_sample(baseline_prompts, tasks)
    cpt_boot = bootstrap_sample(cpt_prompts, tasks)
    v3_boot = bootstrap_sample(v3_prompts, tasks)
    # Compute JS and store
    ...
```

### 4 Supplementary Results

#### 4.1 Token-Level Expert Specialization

**Table 9:** Top-5 tokens for selected experts at Layer 12 (v3 model).

| Expert | Purity | Top-5 Tokens |
|---|---|---|
| E54 | 80% | `the`, `of`, `and`, `to`, `in` |
| E59 | 46% | `AA`, `AT`, `TT`, `GG`, `CC` |
| E78 | 35% | `M`, `L`, `V`, `I`, `F` |
| E97 | 28% | `CD4`, `CD8`, `IL`, `TNF`, `IFN` |
| E100 | 15% | `(`, `)`, `=`, `C`, `O` |

#### 4.2 Per-Task Entropy Analysis

**Table 10:** Shannon entropy of expert distributions across training stages.

| Task | Baseline | CPT | v3 (CPT+SFT) |
|---|---|---|---|
| NL | 3.85 | 4.13 | 4.15 |
| DNA | 2.89 | 2.76 | 2.78 |
| Protein | 2.76 | 2.54 | 2.56 |
| StdHomology | 2.68 | 2.48 | 2.50 |
| RemHomology | 2.71 | 2.52 | 2.54 |
| Cell | 2.95 | 2.88 | 2.90 |
| Mol | 3.02 | 2.95 | 2.97 |
| Structure | 2.82 | 2.65 | 2.67 |

#### 4.3 Supplementary Table S5: Multi-task Generation Evaluation (v5)

**Table 11:** Multi-task generation accuracy across six evaluation categories, 100 samples each, on a held-out split. Keyword-overlap is the standard text-overlap score; Structure additionally reports character-level overlap on the 3Di / DSSP output strings, which are not natural-language tokens. The low Mutation and Literature scores reflect known long-tail and abstract-language difficulties; the dual-head architecture in v5 specifically addresses Structure via the classification heads (Table S6) rather than via generation.

| Category | Keyword score | Char overlap |
|---|---|---|
| Protein | 1.000 | — |
| Mol | 0.980 | — |
| Cell | 0.966 | — |
| Literature | 0.559 | — |
| Mutation | 0.113 | — |
| Structure (gen mode) | 0.000 | 0.259 |

#### 4.4 Supplementary Table S6: Per-residue Classification Head Accuracy (v5)

**Table 12:** Per-residue accuracy of the two classification heads on a 50-protein held-out evaluation, under teacher-forced reference-residue inference. Scores are aggregated by averaging per-protein accuracy. Both heads operate on the same final-hidden-state representation as the generation head; the dual-head architecture trains them jointly with the generation objective ($\alpha = \beta = 0.5$) for 1,350 steps.

| Head | Classes | Chance | Accuracy | N proteins |
|---|---|---|---|---|
| 3Di (Foldseek) | 20 | 5.0% | 78.6% | 19 |
| DSSP (secondary str.) | 8 | 12.5% | 100.0% | 31 |

#### 4.5 Supplementary Table S7: Standard / Remote Homology Updated Numbers

**Table 13:** BioPAWS homology accuracy across all five stages on consistent evaluation samples. Standard uses 1,000 balanced pairs (`protein_pair_short`); Remote uses 500 balanced pairs (`protein_pair_remote`). The v3-to-v4 jump is the headline finding of the v4 recipe change (Alpaca + loss masking + task oversampling).

| Stage | Standard | Remote | BixBench |
|---|---|---|---|
| Gemma-4-Instruct (vocab-extended) | 85.0% | 60.0% | 87.0% |
| v2 (CPT + Bio-SFT) | 100.0% | 56.6% | 91.0% |
| v3 (+20K remote pairs) | 99.95% | 59.5% | 93.7% |
| v4 (Alpaca + loss-mask + oversamp) | 99.5% | 82.0% | 90.4% |
| **v5 (+ dual-head)** | **99.4%** | **82.6%** | **93.7%** |

### 5 Code Availability

All code, training scripts, and analysis notebooks are available at:

`https://github.com/maris205/omnigene4`

#### 5.1 Key Scripts

- `biopaws/cpt/1-prepare_cpt_data_mp.py` – CPT data preparation (multi-process)
- `biopaws/cpt/2-run_cpt.py` – CPT training (8-GPU DDP)
- `biopaws/sft_data/17-train_bio_sft_v3_remote.py` – SFT v3 training
- `biopaws/cpt/40-train_bio_sft_v4.py` – SFT v4 (Alpaca + loss-mask + oversampling)
- `biopaws/cpt/41-train_bio_sft_v5_classifier.py` – SFT v5 (dual-head)
- `biopaws/cpt/18-eval_v3_sft.py` – v3 evaluation
- `biopaws/cpt/42-eval_v4_full.py` – v4 evaluation
- `biopaws/cpt/43-eval_v5_full.py` – v5 evaluation (incl. classification heads)
- `biopaws/cpt/44-merge_v5_to_full.py` – v5 LoRA + heads -> BF16 merge
- `biopaws/cpt/20-collect_moe_activations.py` – Routing collection
- `biopaws/cpt/26-format_matched_routing.py` – Format control
- `biopaws/cpt/27-esm2_remote_headtohead.py` – ESM-2 comparison
- `biopaws/cpt/28-bootstrap_ci.py` – Bootstrap CI

#### 5.2 Data Files

Routing-count matrices and analysis results are in `outputs/moe_analysis/`:

- `moe_counts_baseline.npz` – Baseline routing (50 KB)
- `moe_counts_cpt.npz` – CPT routing (47 KB)
- `moe_counts_v3.npz` – v3 routing (48 KB)
- `format_matched_report.json` – Format control results
- `esm2_remote_2k_eval.json` – ESM-2 comparison results
- `bootstrap_ci_report.json` – Bootstrap CI results
- `bootstrap_samples.npz` – Full bootstrap samples (1,000 iterations)
- `v4_full_eval.json`, `v5_full_eval.json` – v4 / v5 multi-task and classification-head accuracy

##### 5.3 Model Weights

All model artefacts (LoRA adapters, classification heads, merged BF16 weights, tokenizer, chat template) are publicly released on the Hugging Face Hub under the `dnagpt` organization:

- `https://huggingface.co/dnagpt/OmniGene-4-CPT-v2-merged` – Base + CPT only, BF16 ( 50 GB)
- `https://huggingface.co/dnagpt/OmniGene-4-SFT-v3-merged` – CPT + SFT v3, BF16 ( 50 GB)
- `https://huggingface.co/dnagpt/OmniGene-4-SFT-v3-GGUF` – v3 quantized GGUF Q4_K_M ( 16 GB)
- `https://huggingface.co/dnagpt/OmniGene-4-SFT-v4` – v4 LoRA + embedding ( 1.9 GB)
- `https://huggingface.co/dnagpt/OmniGene-4-SFT-v5` – v5 LoRA + embedding + classification heads ( 1.9 GB)
- `https://huggingface.co/dnagpt/OmniGene-4-SFT-v5-merged` – v5 final, BF16 with classification heads ( 52 GB, 11 shards)
